# Single-cell multi-omics enabled discovery of alkaloid biosynthetic pathway genes in the medical plant *Catharanthus roseus*

**DOI:** 10.1101/2022.07.04.498697

**Authors:** Chenxin Li, Joshua C. Wood, Anh Hai Vu, John P. Hamilton, Carlos Eduardo Rodriguez Lopez, Richard M. E. Payne, Delia Ayled Serna Guerrero, Kotaro Yamamoto, Brieanne Vaillancourt, Lorenzo Caputi, Sarah E. O’Connor, C. Robin Buell

## Abstract

Advances in omics technologies now permit generation of highly contiguous genome assemblies, detection of transcripts and metabolites at the level of single cells, and high-resolution determination of gene regulatory features including 3-dimensional chromatin interactions. Using a complementary, multi-omics approach, we interrogated the monoterpene indole alkaloid (MIA) biosynthetic pathway in *Catharanthus roseus*, a source of leading anti-cancer drugs. We identified not only new clusters of genes involved in MIA biosynthesis on the eight *C. roseus* chromosomes but also rampant gene duplication including paralogs of MIA pathway genes. Clustering was not limited to the linear genome and through chromatin interaction data, MIA pathway genes were shown to be present within the same topologically associated domain, permitting identification of a secologanin transporter. Single cell RNA-sequencing revealed exquisite and sequential cell-type specific partitioning of the leaf MIA biosynthetic pathway that, when coupled with a newly developed single cell metabolomics approach, permitted identification of a reductase that yields the bis-indole alkaloid anhydrovinblastine. Last, we revealed cell-type specific expression in the root MIA pathway that is conferred in part by neo- and sub-functionalization of paralogous MIA pathway genes. This study highlights how a suite of omic approaches, including single cell gene expression and metabolomics, can efficiently link sequence with function in complex, specialized metabolic pathways of plants.

## INTRODUCTION

Gene discovery for metabolic pathways in plants has, to date, relied on whole tissue-derived metabolome and transcriptome datasets^1^. Discovering genes from these datasets relies on correlating the expression of genes with the presence of the molecule of interest, along with extensive knowledge of enzymatic biochemical transformations. Occasionally, high quality genome assemblies further facilitate pathway gene discovery by allowing identification of biosynthetic gene clusters, but such spatially clustered groups of pathway genes only occur in limited numbers of plant pathways^2^. Overall, mining of these datasets to identify all of the genes of a complex metabolic pathway typically requires functional screening of large numbers (hundreds or thousands) of candidate genes. These mining approaches are even further limited when genes are not co-regulated or co-localized in the genome. As a consequence, access to the wealth of pharmacologically active molecules encoded in plant genomes has been limited.

In plants, biosynthetic pathways of these complex specialized metabolites (natural products) are localized not only to distinct organs, such as leaves and roots, but also to distinct cell types within these organs and the advent of single cell omics technologies has enormous potential to revolutionize metabolic pathway gene discovery in plants^3–6^. Furthermore, single cell omics comprehensively reveals how metabolic pathways are partitioned across cell types, a central component in understanding specialized metabolite function as well as the precise gene regulation required to confer cell-type specificity of specialized biosynthesis and transport.

The medicinal plant species *Catharanthus roseus* (L.) G. Don produces monoterpene indole alkaloids (MIAs), a natural product family with a wide variety of chemical scaffolds and biological activities^7^. These include the dimeric (bis-indole) MIAs that either demonstrate anti-cancer activity (vinblastine and vincristine)^8^, or are used as a precursor (anhydrovinblastine) for natural and semi-synthetic alkaloids with anti-cancer activity (e.g. vinorelbine^9^). MIA biosynthesis in *C. roseus* shows distinct metabolite profiles across organs, with leaves producing vinblastine and vincristine and roots producing hörhammercine. Over the last 30 years, 38 dedicated MIA pathway genes and several transcription factors involved in jasmonate-induction of the MIA biosynthetic pathway have been discovered using traditional biochemical and co-expression analysis of whole tissue derived omics datasets. Not only does *C. roseus* have enormous economic importance as a producer of anti-cancer drugs^10^, but it has also emerged as a model species to probe the mechanistic basis of localization, transport and regulation of complex specialized metabolic pathways^11^.

Here, we show how a state-of-the-art genome assembly, Hi-C chromosome conformation capture, and single cell transcriptomics datasets empowered novel discoveries in the *C. roseus* MIA biosynthetic pathway. First, we show that a 38-step MIA pathway is sequentially expressed in three distinct cell types in leaves, and second, report on cell-type specific gene expression of the MIA biosynthetic pathway in *C. roseus* roots revealing plasticity in MIA gene expression between organs. Third, we utilized long-range chromosome interaction maps to reveal the 3D organization of MIA biosynthetic gene clusters which contribute to coordinated gene expression in distinct organs and cell types. Fourth, to complement our genomic and transcriptomic datasets, we developed a high throughput, high resolution, semi-quantitative single cell metabolomics profiling method for *C. roseus* leaf cells. Last, to test the power of these emerging omics data, we identified a new intracellular transporter as well as the missing reductase that generates anhydrovinblastine, an important industrial precursor for anti-cancer drugs.

## RESULTS

### A chromosome-scale genome assembly for C. roseus reveals frequent tandem duplication and biosynthetic pathway clustering

Since some specialized metabolic pathways have been demonstrated to be physically clustered in plant genomes, the availability of a high quality, scaffolded genome is essential to accelerate gene discovery^12^. Earlier versions of the *C. roseus* genome assembly were generated using short reads and were highly fragmented^13, 14^. Here, we generated a chromosome-scale, highly contiguous genome assembly of *C. roseus* using state-of-the-art genome sequencing and assembly methods. Using Oxford Nanopore Technologies (ONT) long reads, we generated a draft assembly for *C. roseus* ‘Sunstorm^TM^ Apricot’ and performed error correction using ONT and Illumina whole genome shotgun reads yielding a 575.2 Mb assembly composed of 1,411 contigs with an N50 contig length of 11.3 Mb. Proximity-by-ligation Hi-C sequences were used to scaffold the contigs, resulting in eight pseudochromosomes (Fig. 2a), consistent with the chromosome number of *C. roseus*^15^. To fill gaps within the pseudochromosomes, we capitalized on the ability to redirect in real-time the sequencing of each nanopore using adaptive sampling^16^ by targeting the physical ends of each contig for sequencing, an approach coined ‘adaptive finishing’. We observed 5.5- to 14-fold enrichment of sequence coverage, depending on the length of physical ends that we targeted. Using the adaptive finishing reads along with the bulk ONT genomic reads, we closed 14 gaps in our pseudochromosomes ranging in size from 8-bp to 20.2-kbp. The final (v3) *C. roseus* genome assembly is 572.4 Mb, of which 556.4 Mb is anchored to the eight chromosomes, with an N50 scaffold size of 71.2 Mb (Fig. 2b) representing a significant improvement in contiguity (27.6-fold increase in N50 scaffold length) and 31 Mb genome size increase over v2 of the *C. roseus* genome^14^. Assessment of the v3 *C. roseus* genome using Benchmarking Universal Single-Copy Ortholog (BUSCO) analysis revealed 98.5% BUSCOs indicating a high quality genome assembly.

**Fig. 1:**
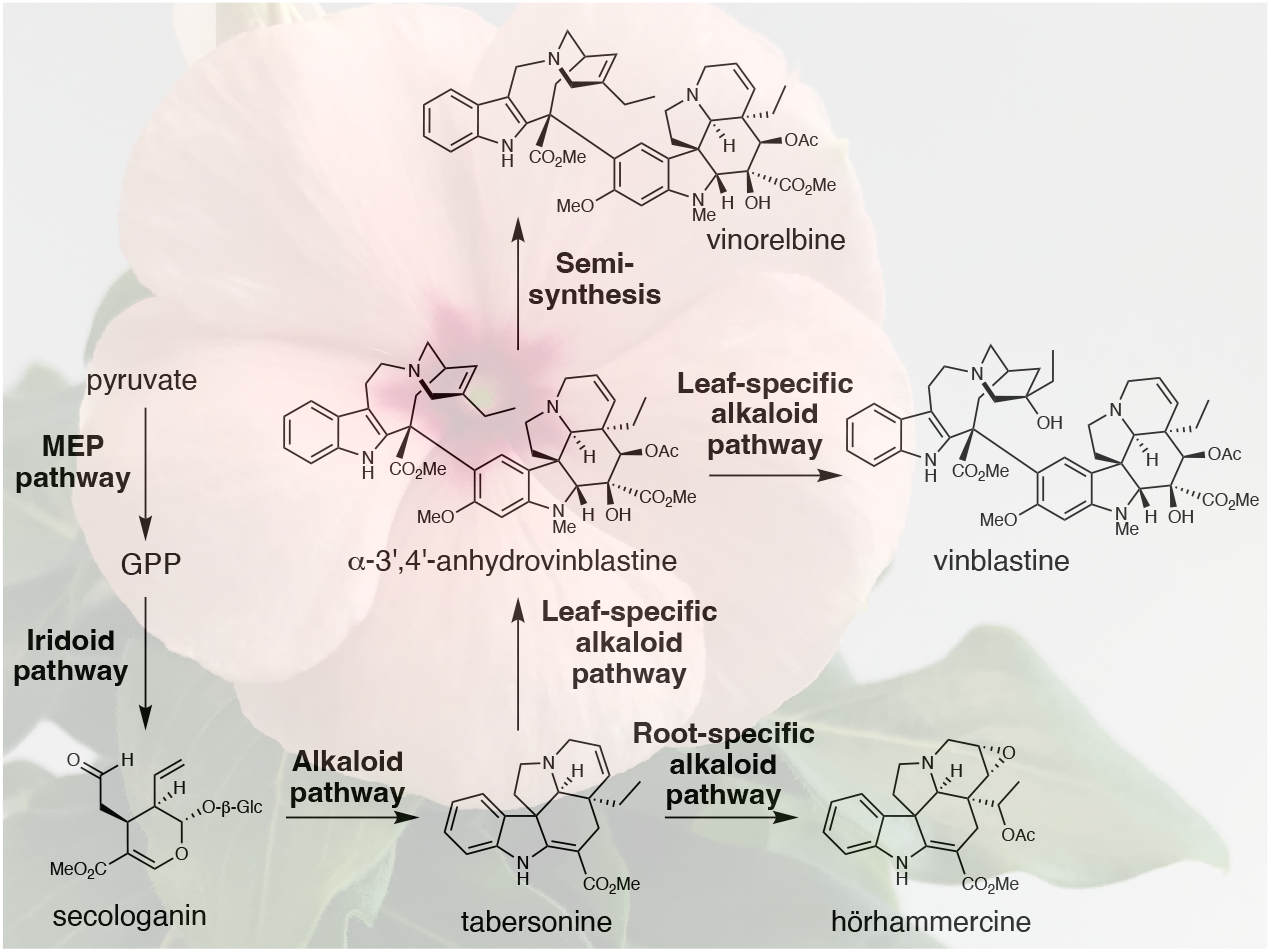
Abbreviated biosynthetic pathway of monoterpene indole alkaloids in *Catharanthus roseus*.

**Fig. 2:**
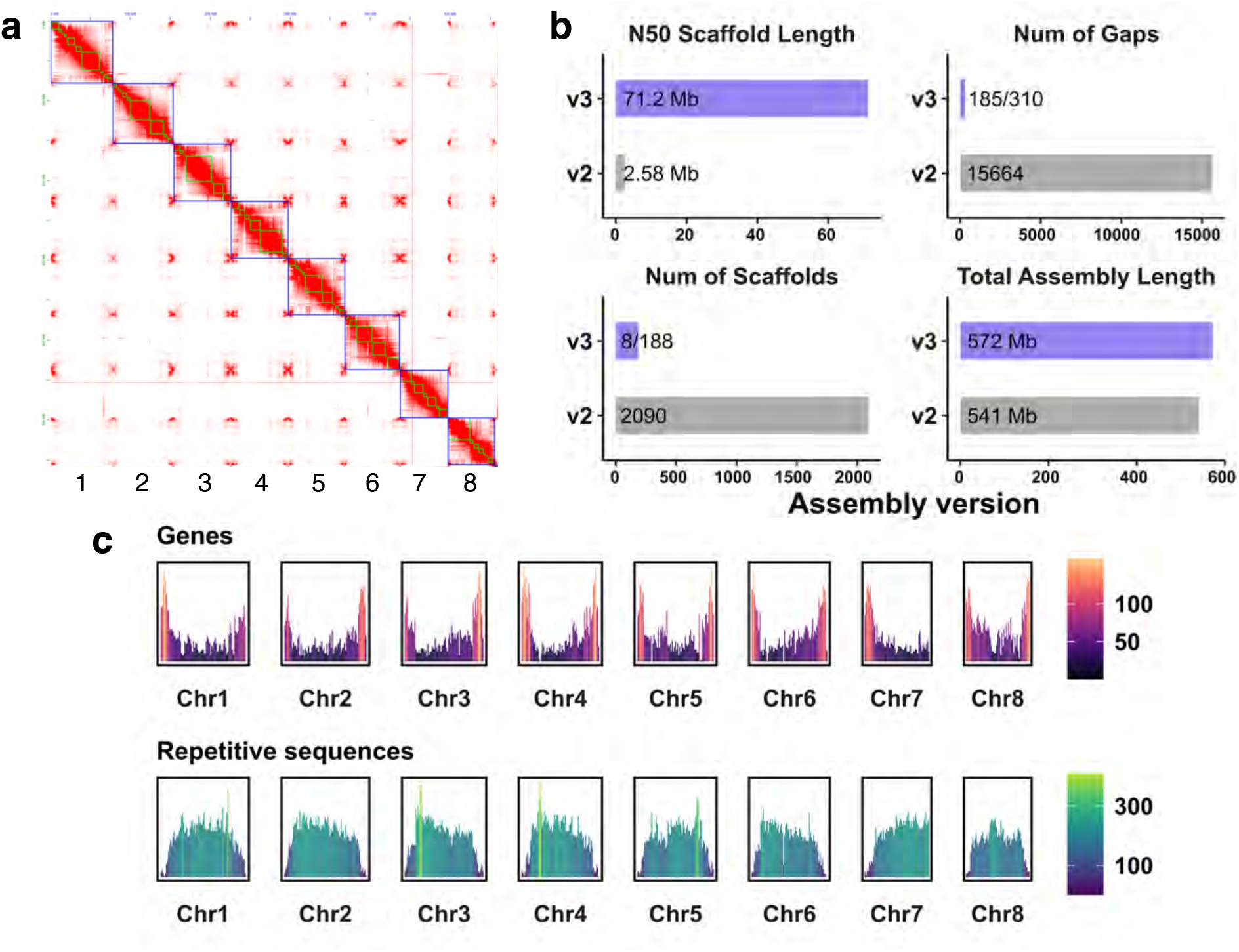
Chromosome-scale assembly and annotation of *Catharanthus roseus*. **a,** Contact map of Hi-C reads revealing the eight chromosomes of *C. roseus*. Blue boxes represent pseudomolecules; green boxes represent contigs of the primary assembly. Red color indicates Hi-C contacts. **b,** Assembly metrics of the v3 vs v2 *C. roseus* genome assembly. For the number of gaps and scaffolds, the v3 numbers represent the pseudochromosomes and the complete assembly (pseudochromosomes plus unanchored scaffolds). **c,** Gene and repetitive sequence density along the eight *C. roseus* chromosomes. y-axis values and color scales represent the number of representative gene models (first row) or repetitive sequences (> 1-kb, second row) in 1-Mb resolution.

To annotate the v3 *C. roseus* genome, we performed *de novo* repeat identification, revealing 70.25% of the genome was repetitive with retroelements being the largest class of transposable elements. Genome-guided transcript assemblies from paired-end mRNA-sequencing (mRNA-seq) reads from diverse tissues (leaf, root, flower, shoots, methyl jasmonate treatment) were used to train an *ab initio* gene finder and generate primary gene models. As empirical transcript evidence provides the highest quality data for gene model construction, we generated 62 million ONT full-length cDNA reads and used these data, along with the mRNA-seq data to refine our gene model structures. The final annotated gene set encompasses 26,347 genes encoding 66,262 gene models with an average of three alternative splice forms per locus, attributable to the deep transcript data used in the annotation. BUSCO analysis of the annotated gene models revealed 96.1% complete BUSCOs, suggestive of high-quality annotation that was confirmed by manual inspection and curation of known MIA pathway genes.

The highly contiguous v3 assembly revealed clusters of MIA pathway genes. Two new clusters were identified, a paralog array containing tetrahydroalstonine synthase 1 (*THAS1)*, *THAS3*, *THAS4* and the *THAS* homologue, heteroyohimbine synthase (*HYS)*, as well as a cluster containing serpentine synthase (*SS*)^17^, a newly identified *SS* paralog with near identical protein sequence, strictosidine glucosidase (*SGD*), and *SGD2*. Gene duplications are major drivers of chemical diversity^18, 19^ and 202 paralogs of 74 MIA genes were identified in the v3 genome, of which, a significant number were locally duplicated. Interestingly, new paralogs of MIA biosynthetic genes were identified within the *SS-SGD-SGD2* cluster, the tabersonine 16-hydroxylase (*T16H2*)*-*16-hydroxytabersonine O-methyltransferase (*16OMT*) cluster, and the strictosidine synthase *(STR*)*-*tryptophan decarboxylase (*TDC*) gene clusters. Notably, a multidrug and toxic compound efflux transporter (*MATE*) was also located in the *STR-TDC* cluster (see also Fig. 3a,b). In summary, the high quality *C. roseus* v3 genome assembly and annotation provides the foundation for accelerated discovery of the final MIA biosynthetic pathway genes and the mechanisms underlying the complex organ and cell-type specific gene regulation of this pathway.

**Fig. 3:**
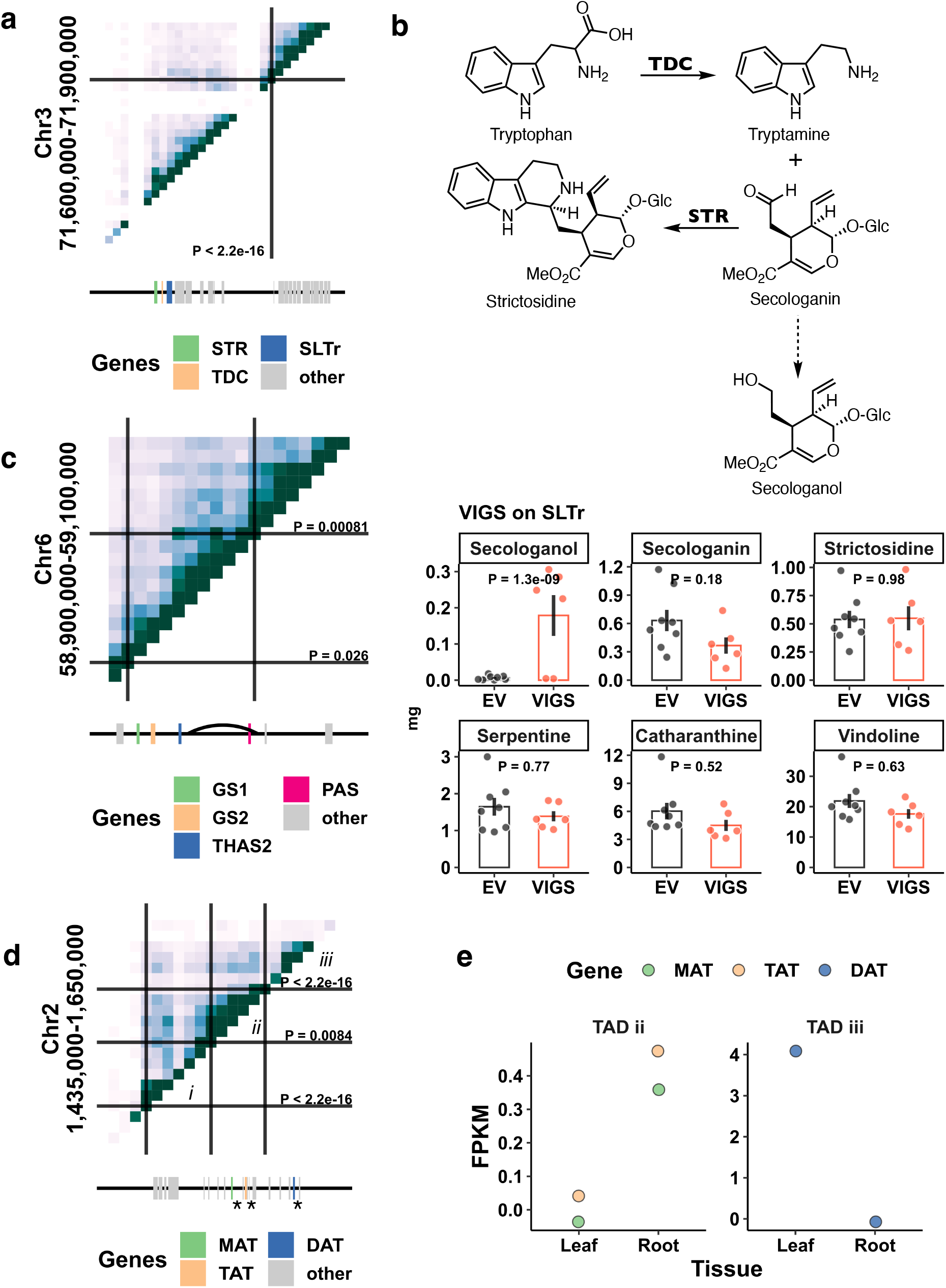
Biosynthetic gene clusters and associated 3-D chromosome features. **a,** Hi-C contact map generated from mature leaves for a gene cluster consisting of *STR*, *TDC* and *SLTr*. **b,** Chemical scheme showing tryptamine, secologanin, strictosidine, and secologanin and virus induced gene silencing (VIGS) for *SLTr*. Bar heights represent means; error bars represent standard errors. Each dot represents a sample. *P* values are based on Tukey tests. EV, empty vector control. **c,** Hi-C contact map for a gene cluster consisting of *GS1*, *GS2*, *THAS2*, and *PAS*. Curve represents the chromosome loop. **d,** Hi-C contact map for a gene cluster consisting of an array of acetyltransferases. i, ii, and iii represent three TADs. Three previously studied biosynthetic genes (*MAT*, *TAT*, and *DAT*) are highlighted by *. **e,** Gene expression profiles for acetyltransferases highlighted in **c**. FPKM: fragments per kilobase exon mapped per million reads. **a, c,** and **d**: Heatmap represents Hi-C contacts at 10-kb resolution, where the darker color represents higher Hi-C contacts. Solid lines and P values represent TAD boundaries detected by HiCKey^21^.

### Biosynthetic gene clusters and associated chromosome conformation features

Unlike biosynthetic gene clusters found in prokaryotic genomes, biosynthetic gene clusters in plant genomes have a loose organization, with unrelated genes or long intergenic spaces separating the biosynthetic genes. Nevertheless, these gene clusters both facilitate gene identification and are believed to play a role in transcriptional regulation^2, 20^. With the recent advent of mapping chromosome conformation features, we now have the ability to probe the location of genes in 3-dimensional (3-D) space. Thus, in addition to searching for biosynthetic gene clusters linearly organized on the chromosome, we can also search for biosynthetic genes that are confined within the same 3-D space.

Using the v3 highly contiguous *C. roseus* genome assembly, we probed chromatin interactions between biosynthetic genes in 3-D space using Hi-C data from mature leaves. HiCKey^21^ was used to detect topologically associated domains (TADs) and HiCCUPS to detect chromosome loops^22^, revealing distinct chromosomal organizations associated with biosynthetic gene clusters. For example, *STR* and *TDC* are physically clustered on chromosome 3 (Fig. 3a). STR and TDC are consecutive steps along the pathway, where TDC catalyzes the formation of tryptamine from tryptophan, and STR catalyzes the condensation between secologanin and tryptamine to form strictosidine. The *MATE* transporter that is located adjacent to *TDC* in the linear genome was also colocated within a TAD with *STR* and *TDC*, as indicated by a high level of long distance contacts detected by Hi-C (Fig. 3a). To test whether this transporter is involved in MIA transport, we performed virus-induced gene silencing (VIGS) of this gene in young *C. roseus* leaves. While the levels of secologanin and downstream alkaloids did not change significantly in response to silencing, we detected a build-up of a compound with *m/z* 391 [M+H]^+^ and *m/z* 413 [M+Na]^+^ in the *SLTr*-silenced tissue but not in empty vector controls (Fig. 3b, P = 1.3×10^-9^, Tukey’s tests). This compound was assigned as secologanol based on co-elution with a standard obtained by chemical reduction of secologanin by NaBH_4_ in methanol. The most likely explanation is that this transporter transports secologanin from the cytosol into the vacuole, where strictosidine synthase is localized. The lack of secologanin transport would result in a build-up of secologanin in the cytosol, where the reactive aldehyde would be reduced to the less toxic secologanol. Thus, we named this MATE transporter *SLTr*. This gene cluster has been observed in the other MIA producers *Gelsemium sempervirens* and *Rhazya stricta*, suggesting that it is conserved across strictosidine producing plants^14^.We also observed that biosynthetic genes interact in 3-D space via chromosome loops as shown with *THAS2* and precondylocarpine acetate synthase (*PAS*)^23, 24^ which are separated by ∼50kb in linear distance (Fig. 3c).

Not all genes in a biosynthetic gene cluster are in the same TAD. For example, an array of locally duplicated acetyltransferases was found on chromosome 2, including three that were previously characterized [minovincinine-19-hydroxy-O-acetyltransferase (*MAT*), tabersonine derivative 19-O-acetyltransferase (*TAT*), and deacetylvindoline O-acetyltransferase (*DAT*)]. This array of acetyltransferases is separated into three TADs with *MAT* and *TAT* within TAD ii, and *DAT* within TAD iii (Fig. 3d)^25–27^. This segregation of acetyltransferases within TADs coincides with organ-level expression patterns; *MAT* and *TAT* are expressed in roots but not in leaves, and *DAT* is expressed in leaves but not in roots (Fig. 3e). These observations suggest chromosome conformation may have regulatory roles in controlling organ-specific biosynthetic gene expression, and consequently, the localization pattern of specialized metabolite production.

### Systematic survey of biosynthetic gene expression at cell-type resolution in leaves using single cell transcriptomics

*In situ* hybridization experiments have established the expression specificity of a subset of the 38 known biosynthetic genes involved in bis-indole alkaloid biosynthesis, where the initial steps are located in inner phloem-associated parenchyma (IPAP) cells, downstream enzymes in the epidermis, and late enzymes located in idioblast cells^28–32^. We performed single cell RNA-sequencing (scRNA-seq) on ∼8-week old *C. roseus* leaves that displayed the typical leaf alkaloid profile and obtained gene expression profiles of ∼12,000 cells and ∼12,000 genes (Fig. 4a). We integrated three independent biological replicates using Seurat^33^ with clustering patterns similar across three replicates. Cell types were assigned using Arabidopsis marker gene orthologs (Fig. 4a); two cell types, IPAP and idioblasts, were inferred using previously studied *C. roseus* biosynthetic genes that show cell type specific expression^28, 29^. In an independent experiment, we profiled gene expression at the single cell level across ∼900 cells using the Drop-seq platform^34^. While fewer cell types were detected using the Drop-seq platform due to lower throughput, expression profiles across the top 3,000 variable genes were highly concordant between the cell types detected by two different platforms. For example, the expression profile of idioblasts detected by the 10x platform is highly concordant with those detected by Drop-seq (*r* > 0.9). Taken together, we inferred that the single cell expression profiles are robust and reproducible across two experimental platforms.

**Fig. 4:**
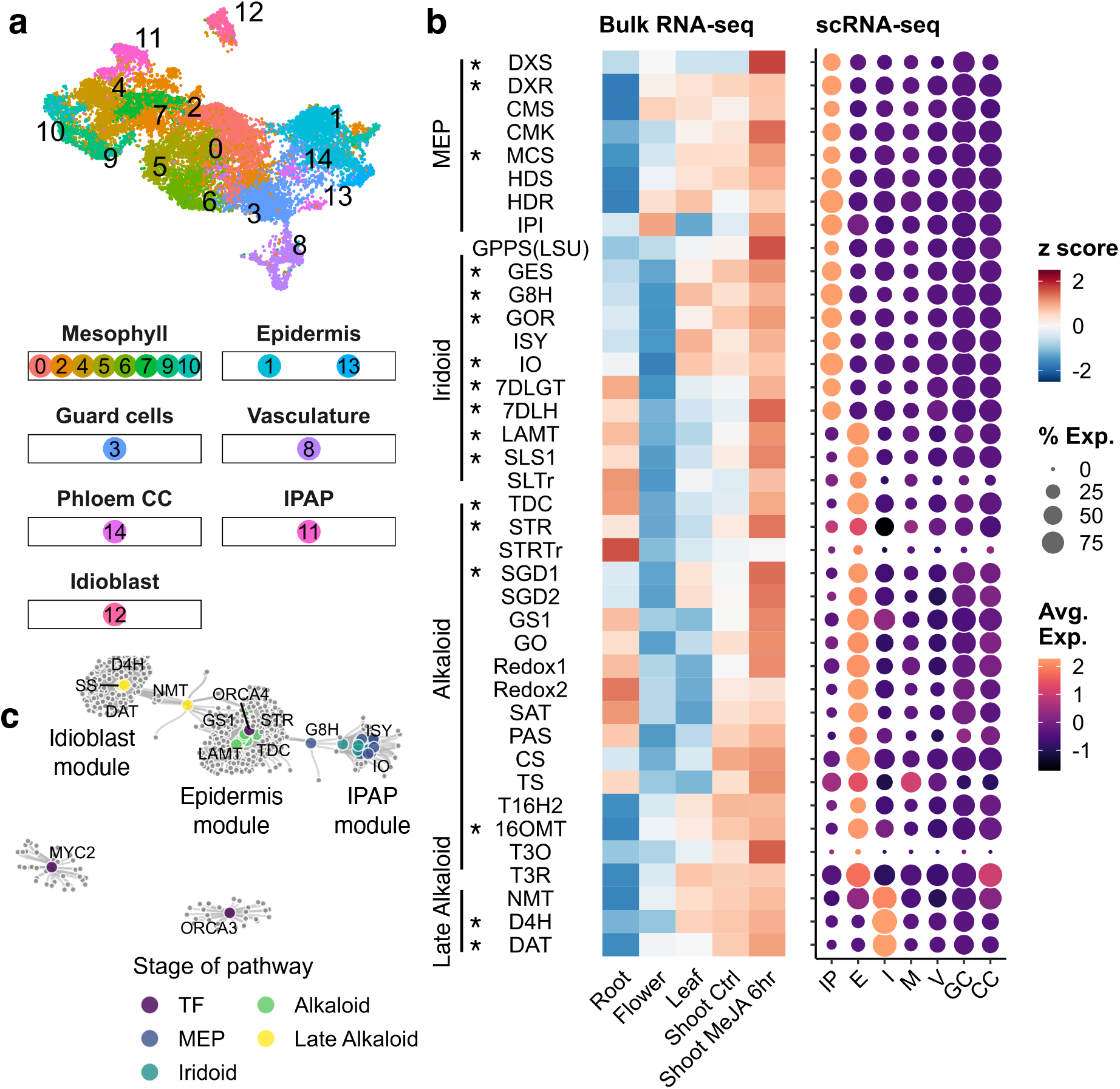
Monoterpene indole alkaloid (MIA) biosynthetic genes are partitioned into three discrete cell types in *Catharanthus roseu*s leaf. **a,** UMAP (uniform manifold approximation and projection) of gene expression in *C. roseus* leaves. **b,** Gene expression heatmap of the MIA biosynthetic pathway for bulk and single cell transcriptomes. Genes are arranged from upstream to downstream. Previously reported cell type specific expression^28–32^ are confirmed and marked with *. **c,** Gene co-expression network for MIA biosynthetic genes using leaf scRNA-seq data. Each node is a gene. Larger size nodes represent previously characterized genes. Edges represent co-expression (FDR < 0.01, see also Methods). IP/ IPAP: internal phloem associated parenchyma; E: epidermis; I; idioblast; M: mesophyll; V: vasculature; GC: stomata guard cells; CC: phloem companion cells.

Co-expression analyses using whole-tissue or organ derived gene expression datasets, i.e., bulk mRNA-seq, has enabled discovery of MIA biosynthetic pathway genes (Fig. 4b). However, whole organ/tissue-derived expression abundances provide an expression estimation that is averaged over all cells in the organ. Thus, if a gene is expressed in a rare cell type such as the idioblast, the power of co-expression analyses will be limited at best. Therefore, we examined scRNA-seq data for biosynthetic gene expression across cell types and found that the pathway is clearly expressed in three specific leaf cell types (Fig. 4b). The improved resolution in cell-type specific expression profiles compared to tissue specific profiles is stark (Fig. 4b). The cell-type specific expression patterns of 16 biosynthetic genes previously characterized by mRNA *in situ* hybridization^28–32^ (Fig. 4b, marked with *) confirmed our cell type resolution of MIA biosynthesis. The MEP and iridoid stages of the pathway are specifically expressed in IPAP whereas the alkaloid segment of the pathway is primarily expressed in the epidermis, and the final known steps of the pathway are exclusively expressed in the idioblasts. Previous work suggested that a heterodimeric GPPS, composed of large subunit (LSU) and small subunit (SSU) is responsible for geranyl pyrophosphate used in MIA biosynthesis^35^. Interestingly, we noted that while the GPPS LSU is specifically expressed in IPAP, the GPPS SSU is expressed across several cell types. Lastly, we found that secologanin transporter (*SLTr)* that is physically clustered with *TDC* and *STR* and co-located within the same TAD as these two biosynthetic pathway genes is specifically expressed in the epidermis (Fig. 3a,b, Fig. 4b), further supporting its involvement in transporting the MIA intermediate secologanin.

We performed gene co-expression analyses producing a network graph for biosynthetic genes as well as previously reported transcription factors. The network self-organized into three main modules, corresponding to IPAP, epidermis, and idioblast (Fig. 4c). The enzyme 3-hydroxy-16-hydroxy-2,3-dihydro-3-hydroxytabersonine *N*-methyltransferase (NMT), which catalyzes a step that bridges the epidermis and idioblast modules on the network diagram, was expressed in both epidermis and idioblast (Fig. 4c). We also note that serpentine synthase (*SS*)^17^ is a member of the idioblast co-expression module. SS catalyzes the formation of serpentine, which has a strong blue autofluorescence and has been previously used as a visual marker for idioblast^17^. The recovery of SS in the idioblast module confirms the robustness of this co-expression analysis. Upon jasmonic acid (JA) elicitation, *MYC2* and *ORCA3* transcription factors are activated, which in turn activate MIA biosynthetic genes^36–39^. However, *MYC2* and *ORCA3* were not part of any modules containing biosynthetic genes (Fig. 4c), suggesting that the regulatory mechanisms in response to JA are distinct from those controlling cell-type specific expression. Lastly, a paralog of *ORCA3*, *ORCA4*^40^, was detected within the epidermis module, suggestive of regulatory roles beyond JA-responsiveness. Taken together, the leaf scRNA-seq dataset is consistent with data obtained from previously established localization methods, and provides accurate and high resolution data for gene discovery. Moreover, the cell-type resolution expression patterns produced co-expression networks that clarified and expanded regulatory relationships.

### High throughput single cell MS data coupled with scRNA-seq

Analysis of metabolic pathways relies on the generation of metabolite-gene expression networks, in which gene expression datasets are analyzed in combination with metabolomic datasets. However, single cell metabolomics (scMS) has lagged behind scRNA-seq due to the intrinsic limitations related to the abundance of the analytes, which is exacerbated by the fact that metabolites cannot be amplified like RNA or DNA. Since the volume of single cells is low (fL to nL range), even when intra-cellular analytes are present at millimolar concentrations, mass spectrometry detection methods require extreme sensitivity. Although progress has been made in the development of scMS approaches^41–43^, few methods have been successfully applied to plant cells. To date, mass spectrometry analyses of individual plant cells have either relied on MS imaging, which is hindered by relatively low spatial resolution, complex sample preparation protocols and low throughput, or live single cell mass spectrometry method (LSC-MS)^44, 45^, which is also highly labor-intensive and not suitable for generation of datasets requiring high throughput. In addition, neither methods utilize chromatographic separation prior to mass spectrometry analysis, greatly limiting accurate structural assignment and absolute quantification of metabolites.

To address these limitations, we designed a process in which a high precision microfluidic cell picking robot was used to collect protoplasts prepared from *C. roseus* leaves from a Sievewell^TM^ device. Protoplasts were then transferred to 96-well plates compatible with a UPLC/MS autosampler. The UPLC/MS method was optimized using available MIA standards. For this study, we collected a total of 672 single cells in seven 96-well plates that were each subjected to UPLC/MS, allowing simultaneous untargeted and targeted metabolomic analysis. As the analysis of all cells was performed over several days, we treated each 96-well plate as an independent experiment, to control for batch-to-batch variation due to experimental and instrumental variables. After close inspection of the selected cells, 86 samples were removed as they either contained two cells or none, reducing the total number of cells included in the analysis to 586. Representative examples of different cells collected in the experiment are shown in Fig. 5a.

**Fig. 5:**
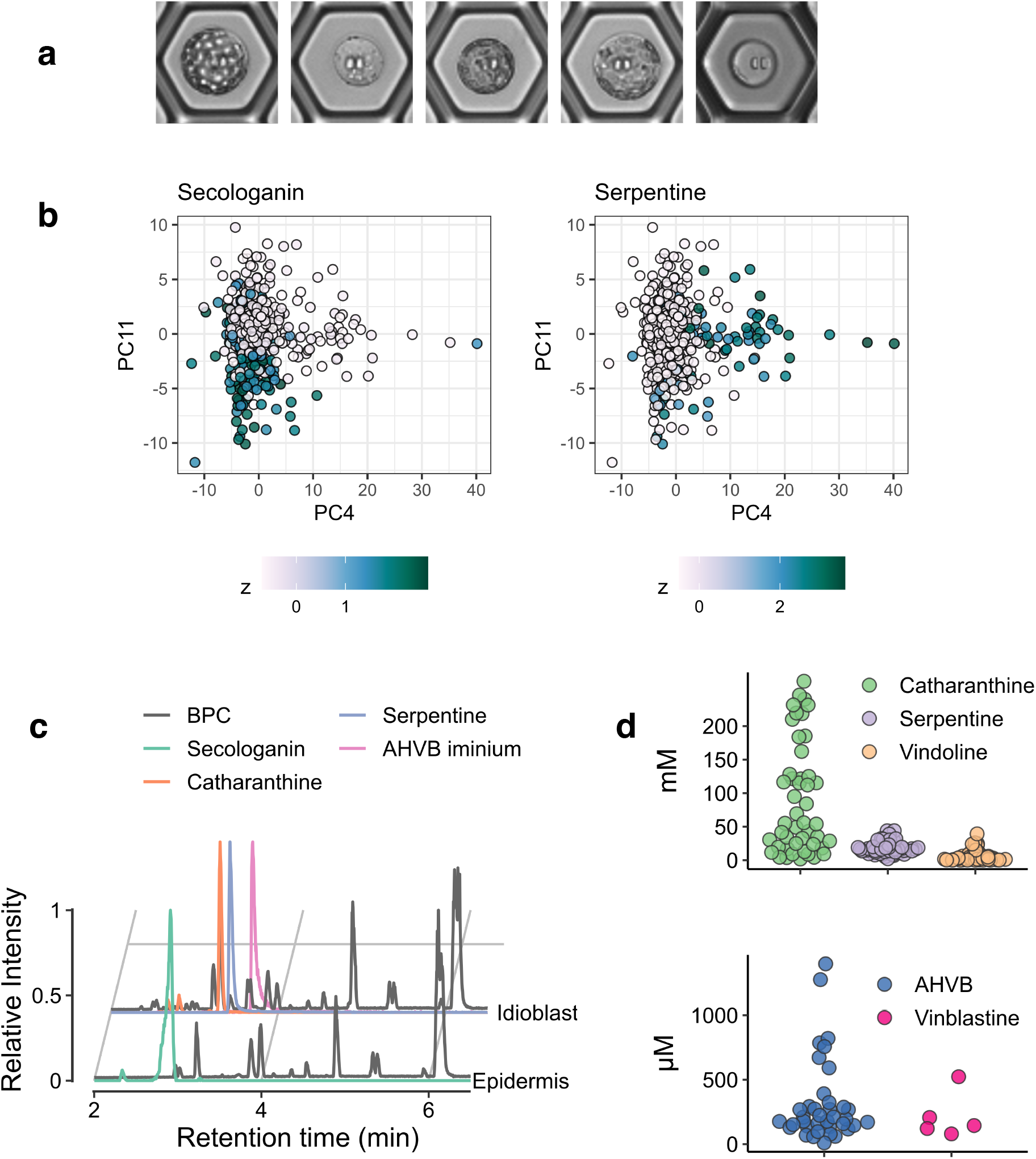
Single cell metabolomic analysis of leaves. **a**, Photos of isolated protoplasts. **b**, Principal component plots colored by z scores of secologanin and serpentine. **c**, LC-MS traces of selected metabolites for idioblast and epidermal cells. BPC: base peak chromatogram of representative epidermis and idioblast cells. **d**, Concentration estimates of selected compounds in single cells.

When using an intensity threshold of 5×10^4^ counts, 34,729 peaks were detected by XCMS. We used CAMERA to group redundant signals (isotopes, adducts, etc.) and kept only the most widely detected peaks, yielding 8,268 representative features. Finally, we excluded the peaks that were not detected in all batches, to a grand total of 933 features. Raw areas were corrected by intensity of the internal standard (ajmaline), log-transformed, and center-scaled by batch to minimize batch-to-batch artifacts. PCA unambiguously separated a group of cells that could be assigned as idioblast cells based on the occurrence of serpentine (*m/z* 349.15467 [M]^+^, C_21_H_20_N_2_O_3_) and vindoline (*m/z* 457.23331 [M+H]^+^, C_25_H_32_N_2_O_6_), which have been previously reported to localize to idioblast cells^44–46^ (Fig. 5b). Additionally, another group of cells, the epidermal cells, could be identified based on the occurrence of the secoiridoid secologanin (*m/z* 389.14422 [M+H]^+^, C_17_H_24_O_10_), which is known to be synthesized in epidermal cells by SLS^47^ (Fig. 5b). Strictosidine, which is formed from secologanin and also synthesized in the epidermis, was observed at low levels in only a few cells, since this compound accumulates in leaves younger than those used here. In spite of our efforts to identify iridoid intermediates that could be markers for IPAP cells, none were detected under these experimental conditions. This was not surprising as these iridoid intermediates do not accumulate at substantial levels and do not ionize efficiently in ESI^+^. Representative chromatograms of an idioblast and of an epidermal cell are shown in Fig. 5c. MS/MS fragmentation experiments, conducted on pooled cells QC samples, allowed unambiguous identification of key MIAs, such as catharanthine (*m/z* 337.19105 [M+H]^+^, C_21_H_24_N_2_O_2_), vindorosine (*m/z* 427.22275 [M+H]^+^, C_24_H_30_N_2_O_5_), vindoline, and serpentine.

External calibration using authentic standards allowed accurate quantification of selected metabolites within the cells. The concentrations of the analytes were corrected for the volume of the cells, calculated from the cell dimensions that were measured during the picking process. To our surprise, the concentration of the major metabolites accumulating in idioblasts was in the millimolar range. Although large differences in concentration were observed in the individual cells, the average catharanthine concentration was 100 mM, which is unexpected since the enzyme that produces this metabolite (catharanthine synthase, CS) is located in the epidermis. We detected only low amounts of catharanthine in a few epidermal cells, and live cell mass spectrometry by Yamamoto et al. also demonstrated that the majority of this MIA accumulates in the idioblast cells^44, 45^. Catharanthine was proposed to be exported from the epidermal cells to the cuticle through the action of the ABC transporter CrTPT2^48^, but although our mass spectrometry method could not be used to measure metabolites on the leaf cuticle, our data clearly show that substantial amounts of catharanthine remain sequestered inside leaf idioblast cells. We hypothesize that catharanthine is rapidly transported from the epidermis to idioblasts. In contrast, the location of most detected MIA, including vindoline, vindorosine and serpentine, correlated perfectly with the cell-type expression of their biosynthetic enzymes^17, 49^.

Our analysis also targeted the bisindole alkaloids (e.g. anhydrovinblastine, vinblastine, and vincristine), which had also been detected in idioblast cells using live single cell mass spectrometry^44^. Anhydrovinblastine (*m/z* 397.21218 [M+2H]^2+^, C_46_H_56_N_4_O_8,_ AHVB) was detected in the micromolar range in almost all of the idioblast cells analyzed and its MS/MS fragmentation confirmed its identity. However, vinblastine (*m/z* 406.21747 [M+2H]^2+^, C_46_H_58_N_4_O_9_) was found only in 5 cells and vincristine was not detected at all. This reflects the low levels in which vinblastine and vincristine accumulate. However, catharanthine and vindoline, the proposed precursors of the bisindole alkaloids, co-occur in the same cell type in which AHVB and vinblastine are present, suggesting that the enzymes involved in bis-indole biosynthesis should also be present in the idioblasts. Since concentrations of catharanthine and vindoline are two orders of magnitude higher than AHVB and vinblastine, it is likely that the coupling reaction leading to the bis-indole alkaloids is a rate limiting step, which could be due to low expression, low specific activity of the coupling enzyme, or that coupling is hindered by intracellular compartmentalization of the two monomers.

The fact that AHVB and vinblastine levels do not correlate with upstream pathway intermediates is likely a major reason why the late stage enzymes that convert vindoline and catharanthine to AHVB and vinblastine have been particularly challenging to elucidate (Fig. 6a). In this coupling reaction, an oxidase activates catharanthine, which then reacts with vindoline to form an iminium dimer. A reductase is then required to reduce this iminium species to form AHVB. Our scMS analysis revealed the presence of a chemical species with *m/z* 396.20436 [M^+^+H]^2+^ and chemical formula C_46_H_54_N_4_O_8_^+^, consistent with this iminium dimer in idioblast cells, along with the monomers and AHVB. The mechanism of enzymatic reduction of the iminium intermediate to AHVB is not known, but recent work in our group has identified a number of medium chain alcohol dehydrogenases in *C. roseus* that can reduce iminium moieties (GS, Redox1, T3R). Therefore, we hypothesized that this reduction of the iminium dimer would be carried out by a medium chain alcohol dehydrogenase localized to idioblast cells. From the sc-RNAseq data, we identified five idioblast-specific medium chain alcohol dehydrogenases (Fig. 6b). These enzymes were heterologously expressed in *E. coli* and assayed with the iminium dimer, which could be generated *in vitro* by incubating catharanthine and vindoline with commercial horseradish peroxidase, which is known to catalyze the initial oxidative coupling. Two enzymes, THAS1 and THAS2, formed AHVB when incubated with the iminium dimer, with specific activity of 10.38 ± 0.27 umol min^-1^ mg^-1^ and 6.26 ± 0.62 umol min^-1^ mg^-1^, respectively (Fig. 6c)^50^. Both of these reductases have been previously biochemically characterized as tetrahydroalstonine synthases, an enzyme that generates tetrahydroalstonine from the reduction of strictosidine aglycone. Notably, however, tetrahydroalstonine levels in leaf are low, which is consistent with an alternative catalytic function for these enzymes in leaf.

**Fig. 6:**
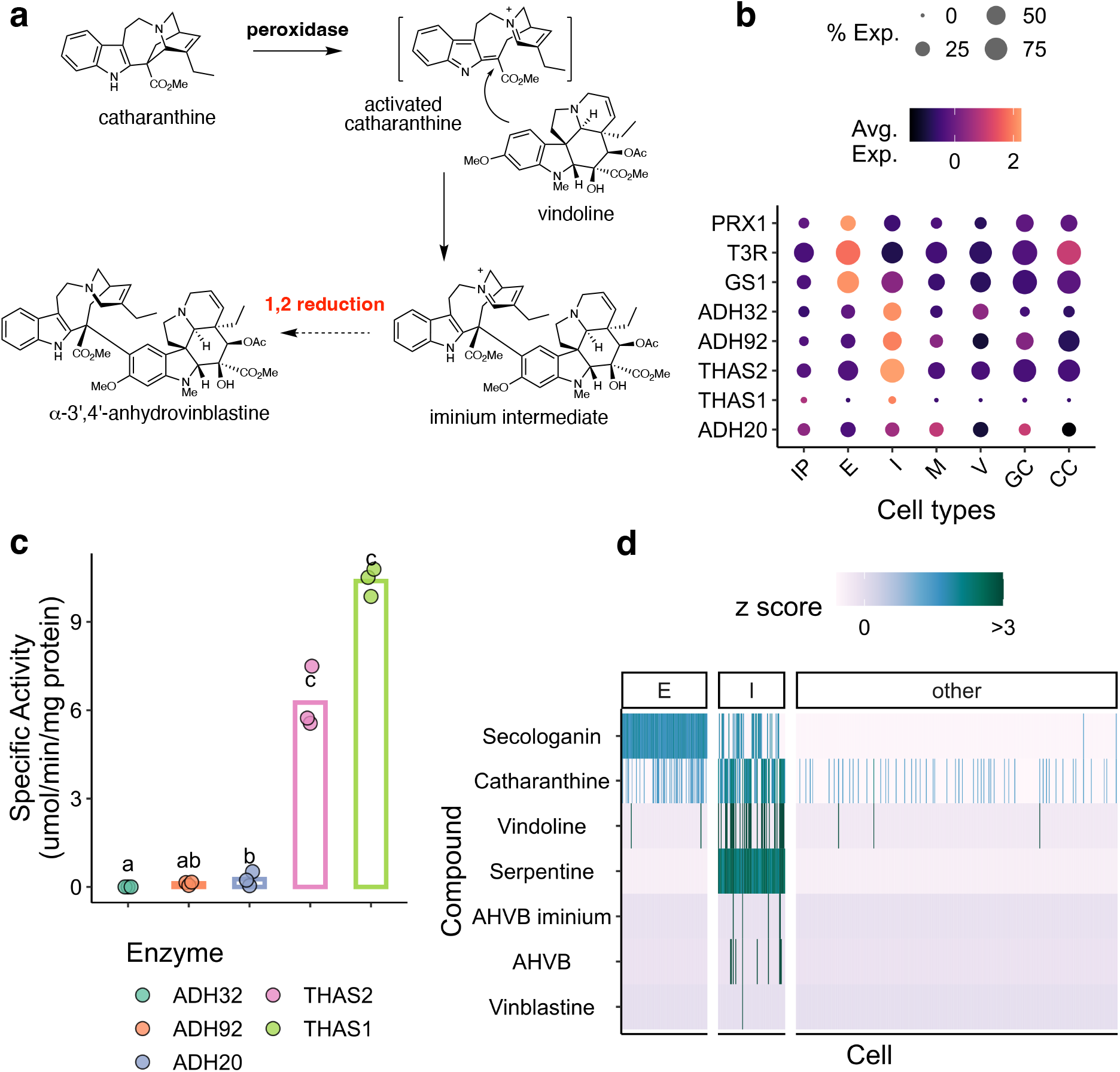
Reduction of an iminium to form anhydrovinblastine. Discovery of ADH20, and comparison of kinetic parameters against THAS2 and other ADHs. **a,** A short chemical scheme showing coupling and reduction. **b,** Expression heatmap at single cell type resolution. **c,** Specific activity of the five idioblast-localized ADH enzymes recombinantly expressed in *E. coli* and tested *in vitro* for activity towards the AHVB iminium. **d,** Heatmap showing cells as columns and compounds as rows from the single cell metabolomics experiment.

A peroxidase, CrPRX1, that can activate catharanthine to form the iminium intermediate, has been previously reported^51^. Surprisingly, this enzyme is selectively expressed in the epidermis (Fig. 6b), in contrast to the localization of vindoline, iminium dimer, and AHVB (Fig. 6d). Notably, the dimerization reaction can be catalyzed by non-specific peroxidases such as horseradish peroxidase, so we hypothesize that CrPRX1 is also a non-selective enzyme that has another function *in planta*. We did not identify any idioblast-specific peroxidase in the leaf scRNA-seq dataset. These omics datasets set the stage for future work, which will focus on functionally characterizing additional classes of oxidases– which are challenging to functionally express in heterologous systems– specifically localized to the idioblast for activity in coupling and oxidation to vinblastine.

### Root single cell transcriptome reveals distinct organization of the MIA biosynthetic pathway

In addition to leaves, we performed scRNA-seq on *C. roseus* roots to compare and contrast MIA biosynthetic gene cell-specific expression in two distinct organs. MIAs are produced in both *C. roseus* leaves and roots, yet while catharanthine and tabersonine are present in both organs, the derivatization of tabersonine diverges in root and leaf. The tabersonine-derived product vindoline, which goes on to form AHVB, vinblastine, and vincristine, is found exclusively in leaves, while tabersonine-derived hörhammercine is found only in roots. The root scRNA-seq dataset captured the expression of ∼2,000 cells from two biological replicates that grouped into 10 clusters and 6 major tissue classes (Fig. 7a). The clustering patterns are highly similar between the two replicates. *MAT* was previously reported by *in situ* hybridization^26^ to be expressed in epidermis and cortex. In the root scRNA-seq data, *MAT* was also found to have dual localization (cluster 4 and cluster 8,). Cluster 4 contained marker genes for both endodermis and cortex (PBL15, AED3)^52^ whereas Cluster 8 contained maker genes for atrichoblast epidermis (TTG2, GL2)^5, 52, 53^. Collectively, these results recapitulate a dual-expressed biosynthetic gene previously characterized by *in situ* hybridization.

**Fig. 7:**
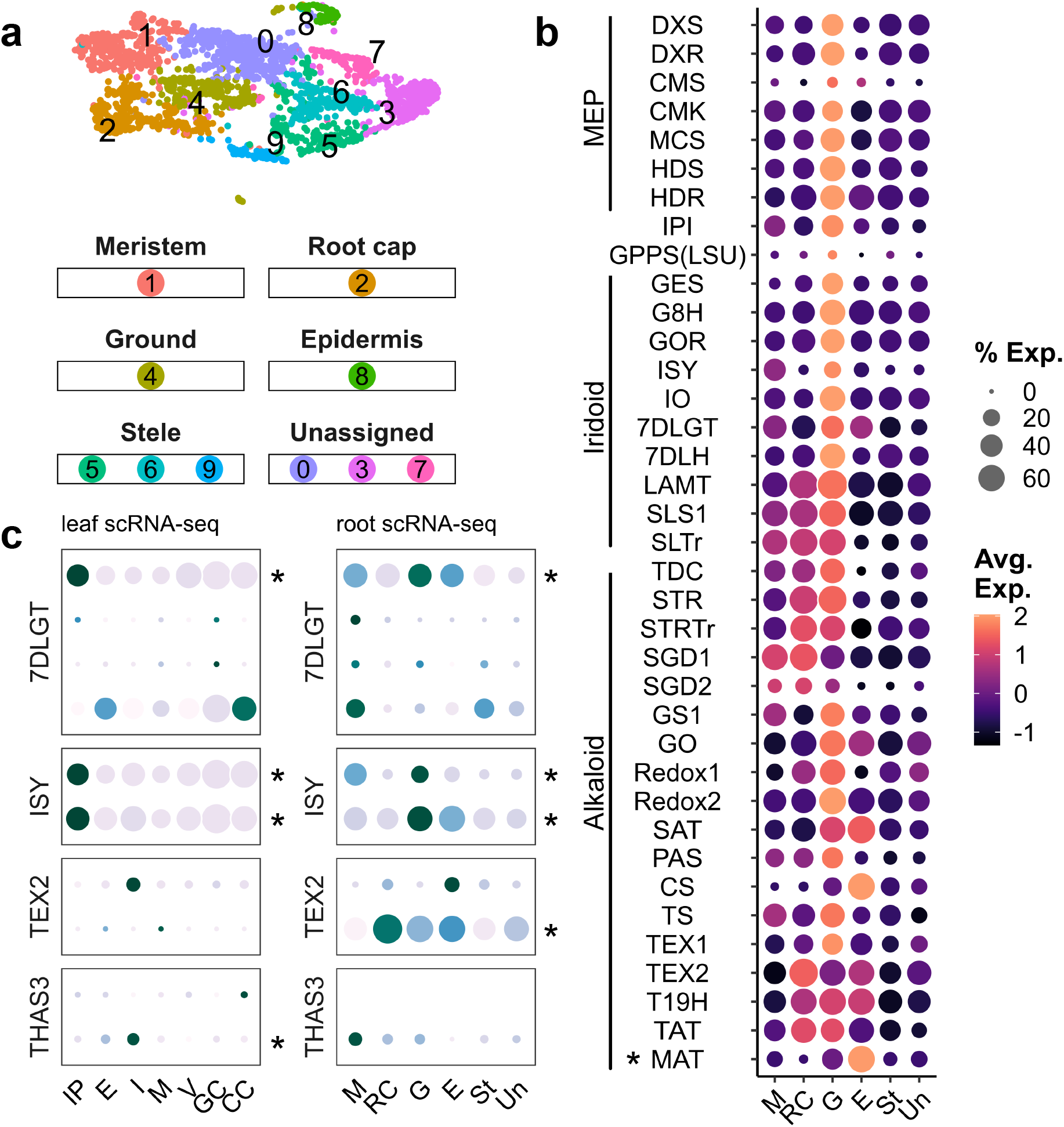
Single cell transcriptome of *C. roseus* roots highlights differential regulation between organs. **a,** UMAP (uniform manifold approximation and projection) for *C. roseus* roots. **b**, MIA biosynthetic gene expression heatmap: Genes are arranged from upstream to downstream. Previously reported cell type specific expression in roots^26^ is marked with *. **c,** Comparison of cell-type specific expression pattern between leaf and root among tandemly duplicated paralogs. Each row is a paralog, organized by gene family. Columns are cell types, grouped by organ. * represents the paralog that is expressed along with the rest of the pathway. Ground tissue includes cortex and endodermis cell types. M: meristem; RC: root cap; G: ground tissue; E: epidermis; St: stele; Un: unassigned.

The spatial organization of the core MIA genes differed between leaf and root, highlighting the plasticity of cell-specific regulation in these two organs. In leaves, the MIA pathway switched from IPAP to epidermal cells at LAMT and from epidermal to idioblast cells at NMT (Fig. 3b), but in roots, the pathway was not partitioned into three discrete cell types (Fig. 7b). Instead, the MEP and iridoid stages are specifically expressed in the ground tissue composed of the cortex and endodermis (Fig. 7b) with expression of the alkaloid stage, while also expressed in ground tissues, exhibiting a more diffused expression pattern (Fig. 7b). Parallel to vindoline biosynthesis in leaves, the late-stage derivatization enzymes that modify tabersonine to hörhammercine are found in a different cell type from the rest of the pathway genes. *TEX*^25^ (tabersonine 6,7-epoxidase), *T19H* (tabersonine 19-hydroxylase)^54^, and *TAT*^55^ (tabersonine derivative 19-O-acetyltransferase) all have detectable expression in the epidermis, along with *MAT*^26^. Our root scRNA-seq experiment used highly developed 8-week old whole root systems which are much more complex than root tips of Arabidopsis seedlings from which the root marker genes for dicots are derived. For example, secondary growth has not occurred in Arabidopsis seedling root tips, while in 8-week old *C. roseus* roots, we clearly observe secondary growth, which may explain the presence of unassigned cell types in our root dataset (Fig. 7a). Despite the highly developed state of the root tissue, we nonetheless observed cell type-specific expression of the MIA pathway primarily in the ground tissue, with the final reaction(s) present in the root epidermis (Fig. 7b).

The v3 assembly resolved multiple tandemly duplicated paralogs, some of which were collapsed or fused in the v2 assembly. To clarify the potential functions of these paralogs, we compared their expression patterns in both leaf and root across cell types. We noticed examples in which a single paralog was preferentially expressed in a given cell type compared to the other paralog(s). For example, a single, collapsed *7DLGT* in v2 was resolved into four separate loci in v3 that displayed cell-type specific expression patterns in leaf and root revealing neo- or sub-functionalization at the expression level (Fig. 7c). We also observed retention of expression patterns among paralogs. Iridoid synthase (*ISY*), well characterized by silencing in leaf, has a tandemly duplicated paralog. Although the *ISY* paralog is expressed in leaf tissue, silencing showed no obvious changes^56^, which can be explained by redundant expression of both paralogs. In the root, both paralogs are expressed in the ground tissue along with the rest of the pathway. However, one of the paralogs is expressed in a higher percentage of cells (Fig. 7c). Lastly, both *TEX2* and *THAS3*^23^ have a tandemly duplicated paralog for which cell type resolution expression patterns highlighted which paralog is likely involved in the biosynthesis of MIAs (marked by *, Fig. 7c).

## CONCLUSION

Prior to the genomics era, few plant specialized metabolic pathways were resolved. However, over the last 15 years, next-generation sequencing technologies allowed the rapid generation of low-cost and high-quality transcriptomic datasets, which, in turn, facilitated gene discovery in a wide range of plant species. Moreover, the availability of draft genome sequences, which have become cost accessible for most plants, enables discovery of physically clustered biosynthetic genes. Here, we report how state-of-the-art omics methods not only accelerate gene discovery through high resolution spatial resolution of gene expression but also how complementary omics data facilitates construction a more holistic view of the genome encapsulating genes, gene regulation in 2-D (linear) and 3-D (chromatin) space, and genic end products, i.e., metabolites.

Generation of a highly contiguous, chromosome-scale genome assembly supported by robust transcript evidence revealed substantial duplication of MIA biosynthetic pathway genes, of which, some are clustered in the linear genome. Chromatin interaction maps revealed 3-D clustering of a subset of MIA pathway genes, some exhibiting organ-level specificity and interactions via chromatin loops, suggestive that TADs, in addition to physical colocalization, can be used to identify genes involved in specialized metabolism. The detection of TADs as well as chromatin loops of physically clustered genes is consistent with the co-regulation hypothesis on the origins of biosynthetic pathway gene clustering as the 3-D chromatin interactions serve key roles in gene regulation^57^. Application of techniques to probe chromatin accessibility such as Assay for Transposase-Accessible Chromatin coupled to high throughput sequencing will reveal *cis*-regulatory sequences including proximal transcription factor binding sites and distal enhancers. The ability to detect cell-type specific gene expression data revealed that the MIA biosynthetic pathway is spatially and sequentially partitioned across discrete cell types in the *C. roseus* leaf permitting construction of cell type-specific co-expression modules for IPAP, epidermis, and idioblast cells. Notably, the cell-type expression profile of MIA biosynthesis genes in leaf and root are substantially different, highlighting the plasticity of the gene expression networks between organs as well as the neo- and sub-functionalization at the gene expression level of MIA biosynthetic pathway paralogs.

Aside from the MIA, multicellular localization patterns of only a few specialized metabolite pathways have been investigated in full. In the morphine biosynthetic pathway, biosynthetic enzymes are synthesized in companion cells, which are then delivered to sieve elements, where the early steps of the pathway take place. Later stage intermediates are then transported from the sieve elements to laticifers where the enzymes involved in the late steps of the morphine pathway are localized which is also the site of morphine and other alkaloid accumulation^58^. Additionally, the localization of the glucosinolate pathway in *Arabidopsis thaliana* has been established^6^. Biosynthetic enzymes for aliphatic glucosinolate biosynthesis appear to be located in xylem parenchyma cells and phloem cells, while indole glucosinolate biosynthetic enzymes are localized to sieve element-neighboring parenchyma cells of the phloem, and glucosinolates are transported to and stored in S cells.

The reasons for the distinct localization of the MIA pathway are not known. The spatial organization of natural products could play an important role in how these metabolites function in defense or signaling. Notably, strictosidine and stricosidine glucosidase, which likely serve an antifeedant role^59^, are located in the epidermis. More derivatized alkaloids, which do not yet have a known ecological role, appear to be derivatized and then stored in the idioblasts, comparable to the role laticifers play in benzylisoquinoline alkaloid biosynthesis. Alternatively, the localization pattern may simply be an accident of evolution, in which the biosynthetic enzymes have evolved from pre-existing enzymes located in these cell types. The partitioning that is observed in MIA biosynthesis does not appear to serve any obvious chemical function, such as separating intermediates that may cross-react.

The high throughput mass spectrometry method developed here showed not only which metabolites co-occurred in distinct cell types, but also allowed us to measure the concentrations of metabolites across a cell population. Notably, although catharanthine is synthesized in the epidermis (as evidenced by the localization of the biosynthetic enzyme CS), this alkaloid co-localizes with vindoline, which is synthesized in idioblasts (as evidenced by the localization of the biosynthetic enzymes D4H and DAT). Therefore, catharanthine must be intercellularly transported from the epidermis to idioblast. Notably, the concentration of catharanthine and vindoline, which dimerize to form anhydrovinblastine (AHVB), were in the high millimolar range. In contrast, the dimerization product AHVB was in the micromolar range, indicating that the coupling step is rate-limiting. This discovery serves as a starting point to design strategies to genetically engineer *C. roseus* plants with higher levels of AHVB.

These state-of-the-art omics datasets provide a foundation for discovery of the remaining genes and regulatory sequences involved in cell-type specific MIA biosynthesis and transport in *C. roseus*. As one-quarter of all pharmaceuticals are derived or have their origins in plants^60^, this study demonstrates the power of single cell multi-omics in natural product gene discovery in plants. We anticipate that application of complementary single cell omics methods will be essential in tapping the wealth of novel chemistries present across the Plant Kingdom.

## MATERIALS AND METHODS

### Genome Sequencing and Assembly

*Catharanthus roseus* cv ‘Sunstorm^TM^ Apricot’ was grown under a 15 hour photoperiod at 22°C and dark-treated for 36 hours prior to harvesting leaves from 17-week old plants. High molecular weight DNA was isolated using a QIAGEN Genomic-tip 500/G after a crude CTAB extraction^61^. ONT libraries were prepared using the ONT SQK-LSK110 kit and sequenced on R9 FLO-MIN106 Rev D flow cells; the latest software available at the run date for each library was used. ONT whole genome shotgun libraries were basecalled using Guppy (5.0.7+2332e8d, https://nanoporetech.com/community) using the high accuracy model (dna_r9.4.1_450bps_hac). Reads less than 10 kb were filtered out using seqtk (v1.3, https://github.com/lh3/seqtk) and remaining reads greater than 10 kb were assembled with Flye (v2.8.3-b1695)^62^ using the parameters -i 0 and --nano-raw. The assembly was polished with two rounds of Racon (v1.4.20)^63^, followed by two rounds of Medaka (v1.4.3, https://github.com/nanoporetech/medaka) using the ‘r941_min_hac_g507’ model, and finally three rounds of Pilon (v1.23)^64^ using Illumina whole genome shotgun reads.

Hi-C libraries were constructed from immature leaf tissue grown under a 15 hour photoperiod with constant 22°C conditions following manufacturer recommendations using the Arima Hi-C 2.0 Kit (Arima Genomics; CRO_AN, CRO_AO). Hi-C libraries were sequenced on a S4 flowcell in paired end mode generating 151 nt reads on an Illumina NovaSeq 6000 (Illumina, San Diego, CA). Contigs less than 10 kb were removed from the assembly using seqtk (v1.3;https://github.com/lh3/seqtk). Pseudochromosomes were constructed using the Juicer (v1.6)^65^ and 3D-DNA (git commit 429ccf4)^66^ pipelines using the Illumina Hi-C sequencing data with default parameters.

To produce the target file for adaptive finishing, 5-kb ends of contigs from the primary assembly were used (full sequences for contigs < 10-kb in size) in the first run while in the second run, 30-kb ends from contigs were used (full sequences for contigs < 60-kb in size). In the second run, half of the channels in the flow cell were set to adaptively sample. Basecalling was performed with Guppy v5.0.16 (nanoporetech.com/community) with the following parameters: --config dna_r9.4.1_450bps_hac.cfg --trim_strategy dna --calib_detect. seqtk v1.3 (github.com/lh3/seqtk) was used to filter reads that were adaptively sampled (not rejected) by the pore. Adaptive finishing reads as well as the bulk ONT genomic reads were used to fill in gaps in the pseudomolecules and unanchored scaffolds using the DENTIST pipeline (v3.0.0)^67^ with the parameters: read-coverage: 90.0, ploidy: 2, max-coverage-self: 3, and join-policy: scaffoldGaps.

### Genome Annotation

A custom repeat library was created using RepeatModeler (v2.0.3)^68^. Putative protein coding genes were removed using ProtExcluder (v1.2)^69^ and Viridiplantae repeats from RepBase^70^ v20150807 were added to create the final custom repeat library. The final genome assembly was hard-masked and soft-masked using the custom repeat library and RepeatMasker (v4.1.2)^71^.

To provide transcript evidence for genome annotation and gene expression abundance estimations, publicly available mRNA-seq libraries were downloaded from NCBI. RNA-seq libraries were processed with Cutadapt (v2.10)^72^ with the parameters: --minimum-length 75 and –quality-cutoff 10. Cleaned reads were aligned to the assembly with HISAT2 (v2.1.0)^73^ with the parameters: --max-intronlen 5000 --rna-strandness ‘RF’ --no-unal --dta; transcript assemblies were generated from the alignments using StringTie2 (v2.2.1)^74^. Full-length cDNA (FL-cDNA) sequences were generated from pooled replicates of young leaf, mature leaf, stem, flower, and root tissue from 16-week old *C. roseus* cv ‘Sunstorm^TM^ Apricot’ plants grown in the greenhouse. RNA was isolated using the Qiagen Rneasy Plant Mini kit followed by mRNA isolation using the Dynabeads mRNA Purification Kit (ThermoFisher Scientific, Waltham, MA, Cat #61011). cDNA libraries were constructed using the ONT SQK-PCB109 kit and sequenced on R9 FLO-MIN106 Rev D flow cells; one library per tissue was constructed and sequenced on a single flow cell. ONT cDNA libraries were basecalled using Guppy (v6.0.6+8a98bbc, nanoporetech.com/community) using the SUP model (dna_r9.4.1_450bps_sup.cfg) and the following parameters: --trim_strategy none -- calib_detect. Base-called reads were processed with Pychopper (v2.5.0, github.com/nanoporetech/pychopper) to identify putative FL-cDNA reads which were then aligned to the genome assembly using Minimap2^75^ (v2.17-r941) with the parameters: -a -x splice -uf -G 5000. Transcript assemblies were generated from the alignments using StringTie2^74^ (v2.2.1).

Initial gene predictions were generated using the BRAKER^76^ (v2.1.5) pipeline using the RNA-seq Stringtie genome-guided alignments as transcript evidence. Gene predictions were refined using the RNA-seq and ONT cDNA transcript assemblies with two rounds of PASA2^77^ (v2.4.1). Monoterpene indole alkaloid biosynthetic pathway genes were manually curated using WebApollo^78^ (v2.6.5). Functional annotation of the gene models was generated by searching the Arabidopsis proteome^79^ (TAIR10), Swiss-Prot plant proteins, and PFAM^80^(v35) and assigning the function from the first informative match.

### Gene expression abundance estimations with bulk mRNA-seq samples

Publicly available *C. roseus* mRNA-seq datasets were downloaded from the NCBI SRA. Reads were cleaned using Cutadapt^72^ (v4.0) to trim adapters and remove low quality sequences. Cleaned reads were aligned to the *C. roseus* genome (v3.0) using HISAT2^73^ (v2.2.1) with a maximum intron size of 5 kb. Fragments per Kilobase Million (FPKM) were generated using Cufflinks^81^ (v2.2.1) with the following parameters; --multi-read-correct, -- max-bundle-frags 999999999, --min-intron-length 20, and --max-intron-length 5000.

### Chromosome conformation capture and analyses methods

*Catharanthus roseus* cv ‘Sunstorm^TM^ Apricot’ leaf tissue collected at the same time as one replicate of the 10x scRNA-seq experiments was used with the Proximo Hi-C kit (Phase Genomics; CRO_AR) to generate Hi-C reads and sequenced on the Illumina NovaSeq 6000 generating paired-end 150 nt reads. Fastq files were processed with Juicer^65^ and Juicer Tools v2.13.07^65^ to produce the .hic file (github.com/aidenlab/juicer/wiki). The inter30.hic output was used for all downstream analyses, which contains chromosome interactions using Q30+ reads. For loops, HiCCUPS (github.com/aidenlab/juicer/wiki/HiCCUPS) was used to detect chromosome loops at 5-kb resolution. For TAD domains, straw (github.com/aidenlab/straw) was used to access the data and write .txt files for each chromosome at 10-kb resolution. HiCkey (github.com/YingruWuGit/HiCKey) was used to detect TAD boundaries; all *p* values are corrected for multiple testing using the false discovery rate method.

### Protoplast Isolation and scRNA-seq Library Generation

Protoplasts were isolated from young leaf tissue of 13-14 week old plants from *C. roseus* cv ‘Sunstorm Apricot’ and used to generate scRNA-seq libraries using the 10x Chromium Controller (10x Genomics) and the Drop-Seq platform. For the 10x Genomics scRNA-seq, leaves were collected, and vacuum infiltrated with enzyme solution (0.4 M mannitol, 20 mM MES, 1.5% [w/v] Cellulase [“Cellulysin”, Sigma, #219466], 0.3% [w/v] Maceroenzyme R-10 [RPI, #M22010], 1 mM Calcium Chloride, 0.1% bovine serum albumin, pH 5.7) for 10 minutes at 400 mbar before being placed into a petri dish and shaken for 1 hour and 45 minutes at 50 RPM. The plates were then shaken at 80-100 RPM for 5 minutes to increase cell recovery. The resulting protoplast solution was filtered through a 40-µm mesh filter into a 50 ml tube with 5ml of storage solution (0.4 M mannitol, 20 mM MES, 1 mM Calcium Chloride, 0.1% bovine serum albumin, pH 5.7) used to rinse the plate and increase cell recovery. Protoplasts were gently pelleted at 150-200 g for 3 minutes at 4 °C and the supernatant removed. The protoplasts were then gently resuspended in storage solution to be counted and used for scRNA-seq library preparation, with additional filtering performed as needed. To generate root single cell expression dataset, ∼5 g of roots from ∼8 week-old plants were used and processed similarly to leaves.

A total of ∼10,700 (leaf), ∼2,600 (leaf), ∼5,800 (leaf), ∼3,400 (root), and ∼4,000 (root) cells were used to generate 10x scRNA-seq libraries. In brief, the protoplast suspensions were loaded into a Chromium microfluidic chip and GEMs generated using the 10x Chromium Controller (10x Genomics); libraries were constructed using the Single Cell 3’ v3.1 Kit (10x Genomics) according to the manufacturer’s instructions. For the Drop-Seq library, scRNA-seq protoplasts were isolated from young leaf tissue from *C. roseus* cv ‘Sunstorm^TM^ Apricot’ and used to generate scRNA-seq libraries following the Drop-Seq method^34^ (mccarrolllab.org/download/905/). In brief, leaves were collected and protoplasts generated as detailed above with the following modifications to the solutions: enzyme solution (0.6 M mannitol, 10 mM MES, 1.5% [w/v] Cellulase [“Cellulysin”, Sigma, #219466], 0.3% [w/v] Maceroenzyme R-10 [RPI, #M22010], 1 mM Calcium Chloride, 0.1% bovine serum albumin, pH 5.7) and storage solution (0.6M, 10 mM MES, 1 mM Calcium Chloride, 0.1% bovine serum albumin, pH 5.7). In total ∼115,000 protoplasts were run through the Drop-Seq protocol to generate the single cell libraries. The 10x Genomics library SCP_AH was sequenced on a NextSeq 500 mid-output flowcell and CRO_AS, CRO_AT, CRO_AW and CRO_AX were sequenced on a NextSeq 2000 P3 flowcell, with all runs sequenced as Read 1 at 28 nt and Read 2 at 91 nt and the index at 8 nt; in accordance with manufacturer recommendations. Drop-Seq libraries CRO_AA and CRO_AB were sequenced on three lanes of an Illumina MiSeq v3, with Read 1 being 25 nt and Read 2 being 100 nt.

### Single Cell Transcriptome Analysis

For 10x Genomics reads, Read 2 was cleaned using Cutadapt^72^ (v4.0) to remove adapters and poly-A tails; cleaned reads were then re-paired with Read 1. Cleaned reads were then aligned to a merged *C. roseus* genome (v3.0) and *C. roseus* chloroplast genome (NC_021423.1) using the STARsolo pipeline of STARsolo (v2.7.10)^82^ with the following parameters: -- alignIntronMax 5000, --soloUMIlen 12, --soloCellFilter EmptyDrops_CR, --soloFeatures GeneFull, --soloMultiMappers EM, --soloType CB_UMI_Simple and --soloCBwhitelist using the latest 10x Genomics whitelist of barcodes.

Drop-Seq reads were processed in accordance with established Drop-Seq processing methods (github.com/broadinstitute/Drop-seq/blob/master/doc/Drop-seq_Alignment_Cookbook.pdf) using DropSeqTools (v2.5.1). Reads were trimmed using Cutadapt^72^ (v4.0) to remove adapters and poly-A tails and aligned to a merged *C. roseus* genome (v3.0) and *C. roseus* chloroplast genome (NC_021423.1) using HISAT2 (v2.2.1)^73^ with the parameters; --dta-cufflinks, --max-intronlen 5000, and --rna-strandness R. The Drop-Seq processing pipeline allows for various filtering approaches in how data is output; a minimum of 100 reads per cell cutoff was used to output our digital expression matrix.

### Cell type clustering

Drop-Seq and 10x Genomics expression matrices were loaded as Seurat objects^33^. Observations were filtered for between 200 and 3000 RNA features and log normalized. Top 3000 variable genes were selected for all runs and integrated using the ‘IntegrateData()’ function from Seurat. UMAP were calculated using the first 20 principal components. We curated a set of epidermis, mesophyll, and vasculature marker genes for *C. roseus* using orthologs^83^ of known markers from Arabidopsis^84, 85^. For root cell markers, we curated markers from Arabidopsis^5, 53, 85, 86^ and *C. roseus*^26^.

### Gene co-expression analyses

Pairwise correlation coefficients between top 3000 variable genes were computed using the ‘cor()’ function in R. A network edge table was produced from the correlation matrix, where each row is a gene pair. Statistical significance was calculated using a t-distribution approximation and adjusted for multiple testing using FDR. Only pairs with FDR < 0.01 were selected for downstream analysis. The network node table was constrained to known MIA biosynthetic enzymes and their 1^st^ degree network neighbors. Graphical representation of the network was produced using the ‘graph_from_data_frame()’ function from igraph^87^, using the filtered edge table and constrained network table as input. Network visualization was done using the ggraph package (ggraph.data-imaginist.com/).

### Leaf protoplast isolation for scMS analysis

Three leaves around 3.5 cm in length were selected. The leaves were cut in 1 mm strips with a sterile surgical blade. After that, the leaf strips were immediately transferred to a Petri dish with 10 mL of digestion medium (2% (w/v) cellulose Onozuka R-10, 0.3% (w/v) macerozyme R-10 and 0.1% (v/v) pectinase dissolved in MM buffer. MM buffer contained 0.4 M mannitol and 20 mM MES, pH 5.7-5.8, adjusted with 1 M KOH. The open Petri dish was put inside a desiccator and 100 mBar vacuum was applied for 15 min to infiltrate the medium into the leaf strips. The vacuum was gently disrupted for 10 seconds after every 1 min. The leaf strips were then incubated in the digestion medium for 2.5 hours at room temperature. After the incubation, the Petri dish was placed on an orbital shaker at around 70 rpm, for 30 min at room temperature to help release the protoplasts. The protoplasts suspension was filtered through a nylon sieve (70 um) to remove larger debris and gently transferred to two 15 mL conical tubes. The protoplast suspension was centrifuged at 70 x g with gentle acceleration/deceleration, for 5 min, at 23 °C to pellet the protoplasts. The supernatant was removed as much as possible and the protoplast pellet of each tube were washed three times by adding 5 mL of MM buffer, swirling gently and centrifuging. Finally, the last pellet from 2 tubes was pooled together and resuspended in 1 mL of MM buffer. The protoplast concentration was determined using a haemocytometer. The final concentration of protoplasts was adjusted to 10^6^ protoplasts in 1 mL.

### Cell picking for scMS analysis

A SIEVEWELL™ chip (ASL) with 90,000 nanowells (50 µm x 50 µm, depth x diameter) was used for single-cell trapping and sorting. The SIEVEWELL™ chip was primed with 100% ethanol and then 1 mL of DPBS was immediately added to the chamber and then discarded through the side ports for washing. This washing step was repeated 2 times. After that, the chip was coated by carefully adding 1 mL 1.5% BSA-DPBS and subsequently discarding the liquid through the side port. Then, MM buffer was added to replace the 1.5% BSA in DPBS solution. Finally, 1 mL of protoplast suspension was carefully added and dispensed in a Z-shape across the well. 1 mL of liquid was then discarded from the side ports.

The SIEVEWELL™ was then mounted on the CellCelector™ Flex (ASL Automated Lab Solutions, Jena) instrument and the cells were visualized using the optical unit, constituted by a fluorescence microscope (Spectra X Lumencor) and a CCD camera (XM-10). Photos in transmitted light were acquired to cover all the chip. Single protoplasts were picked using a 50 µm glass capillary and dispensed into 96-well plates containing 5 µL of 0.1% formic acid. Pictures of the nanowells before (with single-cell) and after picking (without single-cell) were recorded. Cells were dried overnight and frozen at −20°C until the analysis.

### scMS method

The UPLC/MS analysis was performed on a Vanquish (Thermo Scientific) system coupled to a Q-Exactive Plus (Thermo Scientific) orbitrap mass spectrometer. For metabolite separation, a Phenomenex Kinetex® XB-C18 100 Å column (50 x 1.0 mm, 2.6 µm) was used at a temperature of 40 °C. The binary mobile phases were 0.1% HCOOH in MilliQ water (aqueous phase) (A) and ACN (B). The gradient elution started with 99% of the aqueous phase, and increased with ACN phase to 70% in 5.5 minutes, followed by an increase to 100% in the next 0.1 min. The percentage of ACN was held at 100% for 2 minutes before switching back to 99% of the water phase in 0.1 min. Finally, ACN was kept at 1% for 2.5 minutes in order to condition the column for the next injection. Total time for chromatographic separation was 12 minutes. 4 µL of standards or samples were injected into the column via the autosampler. The flow rate of the mobile phase was kept constantly at 0.3 mL/min during the chromatographic separation. Both samples and standard solutions were kept at 10°C in the sample tray. The needle in the autosampler was washed using a mixture of methanol and MilliQ water (1:1, v:v) for 20 seconds after draw and at a speed of 50 uL/s.

The mass spectrometer was equipped with a heated electrospray ionization (HESI) source. The mass spectrometer was calibrated using the Pierce positive and negative ion mass calibration solution (Thermo Scientific). The operating parameters of HESI are based on the UPLC flow rate of 300 µL/min using source auto default: sheath gas flow rate at 48; auxiliary gas flow rate at 11; sweep gas flow rate at 1; spray voltage +3500 V; capillary temperature at 250°C; auxiliary gas heater temperature at 300°C; and S-lens RF level at 50. Acquisition was performed in full scan MS mode (resolution 70000-FWHM at 200 Da) in positive mode over the mass range *m/z* from 120 to 1000. The full MS/dd-MS2 (full-scan and data-dependent MS/MS mode) was used to simultaneously record the MS/MS (fragmentation) and the spectra for the precursors of QC pooled samples. The full MS/dd-MS2 (that included target analytes) was also used for QC pooled sample to confirm fragments of the selected precursors. The dd-MS2 was set up with the following parameters: resolution 35000 FWHM; mass isolation window 0.4 Da; maximum and minimum automatic gain control (AGC) target 8 × 10^3^ and 5 × 10^3^, respectively; normalized collision energy (NCE) was set at 3 levels 15%, 30% and 45%; spectrum data format was centroid. All the parameters of the UHPLC-HRMS system were controlled through Thermo Scientific Xcalibur software version 4.3.73.11 (Thermo Scientific). Chromatography and MS response were optimized using several reference compounds.

### Preparation of cells and quality control (QC) samples

Before the analysis, the single cells were resuspended with 12 µL of 0.1% formic acid containing 10 nM of ajmaline as an internal standard. For pooled QC samples, 2 µL of sample from each well was taken and pooled together. For QC, 20 µL of *C. roseus* leaf protoplasts were extracted with 500 µL of pure MeOH. After sonication (10 min) and vortexing, the protoplast extract was filtered, diluted 200-fold with 0.1% formic acid containing 10 nM of ajmaline and used as a QC run. For QC total, we used a methanolic extract of one of the leaf strips used for making protoplasts. The tissue was ground to a fine powder using a Tissuelyser II (Qiagen). Metabolites were extracted from the powdered leaf sample with 300 µL of pure MeOH. After vortexing and sonication for 10 min, the leaf extracts were filtered, and 5 µL aliquots were placed into Eppendorf tubes and dried under vacuum. Before analysis, one Eppendorf tube was taken out, resuspended in 1 mL of the extraction solution (0.1% formic acid containing 10 nM of ajmaline), sonicated for 10 min, filtered and used as a QC total.

### Preparation of standard solutions and calibration curves

Catharanthine (Sigma Aldrich), vindoline (Chemodex), serpentine hydrogen tartrate (Sequoia Research Products), anhydrovinblastine disulfate (Toronto Research Chemicals) and vinblastine sulfate (Sigma Aldrich) were dissolved in pure MeOH at a concentration of 10 µM. Serial dilutions (n=15) were made between 0.1 and 1000 nM and analyzed by UPLC/MS. The extracted peak areas were used to calculate linear regression curves.

### XCMS analysis and statistical analysis

Peak detection was performed using XCMS^88^ centWave^89^ algorithm with a prefilter intensity threshold of 5×10^4^ counts, a maximum deviation of 3 ppm, a signal to noise ratio greater than 5, peak width from 5 to 30 seconds, integrating on the real data. Peaks were grouped using a density approach with a bandwidth of 1, retention time was corrected with a locally estimated scatterplot smoothing (loess) and a symmetric fit (Tukey’s biweight function), and peaks were re-grouped after correction. Finally, gap filling was performed by integrating raw data in each peak group region. We used CAMERA^90^ to group redundant features (isotopes, adducts and in-source fragments) taking the injections of the QCs of the pooled samples as representative runs. Peaks were grouped within a window of 50% of the full width at half maximum (FWHM); isotopes were detected for single and doubly charged species with an error of 1 ppm; pseudospectra were grouped by within sample correlation of extracted ion chromatograms, with a correlation threshold of 0.85 and a p-value threshold of 0.05; and finally adducts were determined for single cluster ions with 1 ppm error.

Only one representative feature for each correlation group was selected, with preference given to the peaks detected in the largest number of single-cell samples, and breaking ties by total sum of intensities. To reduce variation due to injection, we scaled raw areas by the recovery of the internal standard (ajmaline) in each run. Artifacts were removed by keeping only features that were detected in all injection batches, and the ajmaline-corrected, log-transformed areas were centered and scaled in a per-batch basis, to minimize the effect of batch-to-batch variation. Principal Component Analysis was performed on this matrix using the *base* library of the R programming language (v4.1.3).

We assigned the identities of detected features by searching the exact mass within the limits of the feature *m/z* as detected by XCMS, and the retention time within 5 seconds of the experimental elution time of the standard, and we manually verified the assignations by comparing the QC runs against an injection of a mix of standards that was performed at the beginning and end of each batch.

### VIGS of MATE transporter

A 516 bp fragment of the *SLTr* transporter was amplified from *C. roseus* Sunstorm Apricot leaves cDNA and cloned into the pTRV2u vector as previously described^47^. *Agrobacterium tumefaciens* GV3101 was transformed with the construct by electroporation. Agrobacterium GV3101 strains containing pTRV1, pTRV2u-empty vector, pTRV2u-MgChl (for silencing magnesium chelatase) and pTRV2u-SLTr, were grown overnight in 5 mL LB supplemented with rifampicin, gentamycin and kanamycin at 28 °C. Cultures were pelleted at 3,000 *g*, resuspended in inoculation solution (10 mM MES, 10 mM MgCl_2_, 200 µM acetosyringone) to an OD600 of 0.7, and incubated at 28 °C for 2 h. Transformants were confirmed by PCR using the gene-specific primers used to amplify the gene fragment. Strains containing pTRV2 constructs were mixed 1:1 with pTRV1 culture and this mixture was used to inoculate plants by the pinch wounding method. Plants (8–12, 4 weeks old) were inoculated for each construct, and the plants were grown at 25 °C in a 12 h photoperiod. The pTRV2u was used as a negative (empty vector) control, and the pTRV2-MgChl plants were used as a visual marker of the silencing response, with bleaching of the leaves occurring 21–25 days post-inoculation. On silencing, the plant material was harvested, ground in a TissueLyserII (Qiagen) and stored at – 80 °C before analysis by quantitative PCR (qPCR) and LC/MS. qPCR was performed as reported previously^47^.

For LC/MS of the VIGS tissue, 20 mg of frozen powder were extracted in 600 µL of methanol containing 40 µM caffeine as an internal standard and incubated at 60 °C for 2 h. After a 30 min centrifugation step at 5,000 *g*, an aliquot of the supernatant (25 µL) was mixed with an equal volume of water and analyzed on a Shimadzu IT-TOF instrument. Chromatography was performed on a Phenomenex Kinetex 5 µm C18 100 Å (100 × 2.10 mm × 5 µm) column and the binary solvent system consisted of acetonitrile (ACN) and 0.1% formic acid in water. The elution programme was as follows: 1 min isocratic at 12% ACN, 3.5 min gradient up to 25% ACN, 2.5 min gradient up to 50% ACN, 1 min gradient up to 100% ACN, 6 min isocratic at 100% ACN, 1 min gradient down to 12% ACN and 2.5 min isocratic at 12% ACN. Peak areas were calculated using the LCMS solutions software and normalized by leaf mass (fresh weight) and the peak area of the internal standard.

### Protein expression and activity assays

Cloning of THAS1 and THS2 was described in Stavrinides et al.^23^. ADH20 (KU865330.1), ADH32 (AYE56096.1), and ADH92 (XXXXX) genes were amplified from *C. roseus* ‘Sunstorm Apricot’ leaves cDNA. The PCR products were purified from agarose gel, ligated into the BamHI and KpnI restriction sites of the pOPINF vector^91^ using the In-Fusion kit (Clontech Takara) and transformed into chemically competent *E. coli* Top10 cells. Recombinant colonies were selected on LB agar plates supplemented with carbenicillin (100 µg/mL). Positive clones were identified by colony PCR using T7_Fwd and pOPIN_Rev primers. Plasmids were isolated from positive colonies grown overnight. Identities of the inserted sequences were confirmed by Sanger sequencing. Chemically competent SoluBL21 *E. coli* cells (Amsbio) were transformed by heat shock at 42 °C. Transformed cells were selected on LB agar plates supplemented with carbenicillin (100 µg/mL). Single colonies were used to inoculate starter cultures in 10 mL of 2 x YT medium supplemented with carbenicillin (100 µg/mL) that were grown overnight at 37 °C. Starter culture (1 mL) was used to inoculate 100 mL of 2 x YT medium containing the antibiotic. The cultures were incubated at 37 °C until OD600 reached 0.6 and then transferred to 16 °C for 30 min before induction of protein expression by addition of IPTG (0.2 mM). Protein expression was carried out for 16 h. Cells were harvested by centrifugation and re-suspended in 10 mL of Buffer A (50 mM Tris-HCl pH 8, 50 mM glycine, 500 mM NaCl, 5% glycerol, 20 mM imidazole,) with EDTA-free protease inhibitors (Roche Diagnostics Ltd.). Cells were lysed by sonication for 2 minutes on ice. Cell debris was removed by centrifugation at 35,000 *g* for 20 min. Ni-NTA resin (200 uL, Qiagen) was added to each sample and the samples were incubated at 4 °C for 1 h. The Ni-NTA beads were sedimented by centrifugation at 1000 rpm for 1 min and washed three times with 10 mL of Buffer A. The enzymes were step-eluted using 600 µL of Buffer B (50 mM Tris-HCl pH 8, 50 mM glycine, 500 mM NaCl, 5% glycerol, 500 mM imidazole) and dialyzed in Buffer C (25 mM HEPES pH 7.5, 150 mM NaCl). Enzymes were concentrated and stored at −20 °C prior to in vitro assays.

To assay the activity of the ADHs, the substrate (AHVB iminium) first had to be generated *in vitro*. For this purpose, 500 µL reactions were assembled in 50 mM MES buffer pH 6.5. Each reaction contained 100 µM vindoline, 600 µM catharanthine, 0.002% hydrogen peroxide and 22.5 U of Horseradish peroxidase (Sigma 77332). The reactions were incubated at 30 °C for 45 min, after which the sample was divided into aliquots of 70 µL each. To each aliquot, NADPH was added to a concentration of 200 µM and the ADHs to a concentration of 1 µM. The final volume was 100 µL. The reactions were incubated for 2 h and 3 µL were taken at different time points to monitor the progression of the reactions. The 3 µL samples were quenched in 97 µL of MeOH, filtered and analyzed by UPLC/MS on a Thermo Ultimate 3000 chromatographic system coupled to a Bruker Impact II mass spectrometer. Separation was performed on a Phenomenex Kinetex 2.7 µm C18 100 Å (100 × 2.10 mm × 2.7 µm) column and the binary solvent system consisted of ACN and 0.1% formic acid in water. The elution program was as follows: time 0 to 1 min, 10% ACN; linear gradient to 30% ACN in 5 min; column wash at 100% ACN for 1.5 min, then re-equilibration to 10% ACN for 2.5 min. Flow rate was 600 uL/min. Data were analyzed using the Bruker Data Analysis software. Quantification of the AHVB produced during the reactions was performed using external calibration curves and used to calculate the specific activity of the enzymes. The experiments were performed in triplicate.

## ACKNOWLEDGEMENTS

Funding for this project was provided by the Georgia Research Alliance (C.R.B), University of Georgia (C.R.B.), Michigan State University (C.R.B.), the European Research Council (788301) (S.E.O.), Max Planck Society (S.E.O.) and KAKENHI (20J00973) (K.Y.). Sequencing was performed at the Georgia Genomics and Bioinformatics Core at the University of Georgia and the Research Technology Support Facility at Michigan State University. Library preparation and sequencing of library CRO_AR was performed at Phase Genomics (Seattle, WA). We gratefully acknowledge K. Gase and K. Luck for technical assistance with cloning. We gratefully acknowledge S. Freyberg and J. Langner from ALS Automated Lab Solutions (Jena) for assistance with cell picking for single cell mass spectrometry.

## AUTHOR CONTRIBUTIONS

C.L. generated scRNA-seq and adaptive sampling data. J.C.W. generated genome, Hi-C, scRNA-seq and full-length cDNA sequence data. J.P.H. generated the genome assembly and annotation. A.V., L.C. and D.A.S.G. developed and collected all scMS data. L.C., K.Y. and R.M.E.P. assayed all enzymes and transporters. C.L., J.C.W., J.P.H. B.V., L.C. and C.E.R.L performed data analyses. C.L., J.C.W., J.P.H., C.E.R.L, B.V., L.C., S.E.O., C.R.B. wrote the manuscript. S.E.C. and C.R.B. conceived the study. All authors approved the manuscript.

## Notes

### Competing Interest Statement

The authors have declared no competing interest.

## REFERENCES

1. Zhao, K. & Rhee, S. Y. Omics-guided metabolic pathway discovery in plants: Resources, approaches, and opportunities. Curr. Opin. Plant Biol. 67, 102222 (2022).

2. Polturak, G. & Osbourn, A. The emerging role of biosynthetic gene clusters in plant defense and plant interactions. PLoS Pathog. 17, e1009698 (2021).

3. Kang, M., Choi, Y., Kim, H. & Kim, S.-G. Single-cell RNA-sequencing of *Nicotiana attenuata* corolla cells reveals the biosynthetic pathway of a floral scent. New Phytol. (2022) doi:10.1111/nph.17992.

4. Cuperus, J. T. Single-cell genomics in plants: current state, future directions, and hurdles to overcome. Plant Physiol. 188, 749–755 (2022).

5. Shulse, C. N. et al. High-Throughput Single-Cell Transcriptome Profiling of Plant Cell Types. Cell Rep. 27, 2241–2247.e4 (2019).

6. Tenorio Berrío, R., et al. Single-cell transcriptomics sheds light on the identity and metabolism of developing leaf cells. Plant Physiol. 188, 898–918 (2022).

7. O’Connor, S. E. & Maresh, J. J. Chemistry and biology of monoterpene indole alkaloid biosynthesis. Nat. Prod. Rep. 23, 532–547 (2006).

8. Martino, E. et al. Vinca alkaloids and analogues as anti-cancer agents: Looking back, peering ahead. Bioorg. Med. Chem. Lett. 28, 2816–2826 (2018).

9. o Synthesis and biological evaluation of vinca alkaloids and phomopsin hybrids. J. Med. Chem. 52, 134–142 (2009).

10. Zhu, J., Wang, M., Wen, W. & Yu, R. Biosynthesis and regulation of terpenoid indole alkaloids in *Catharanthus roseus*. Pharmacogn. Rev. 9, 24–28 (2015).

11. Facchini, P. J. & De Luca, V. Opium poppy and Madagascar periwinkle: model non-model systems to investigate alkaloid biosynthesis in plants. Plant J. 54, 763–784 (2008).

12. Nutzmann, H. W., Huang, A. & Osbourn, A. Plant metabolic clusters - from genetics to genomics. New Phytol. 211, 771–789 (2016).

13. Kellner, F. et al. Genome guided investigation of plant natural product biosynthesis. The Plant Journal 82, 680–692 (2015).

14. Franke, J. et al. Gene Discovery in Gelsemium Highlights Conserved Gene Clusters in Monoterpene Indole Alkaloid Biosynthesis. Chembiochem 20, 83–87 (2019).

15. Guimarães, G. et al. Cytogenetic characterization and genome size of the medicinal plant *Catharanthus roseus* (L.) G. Don. AoB Plants 2012, ls002 (2012).

16. Loose, M., Malla, S. & Stout, M. Real-time selective sequencing using nanopore technology. Nat. Methods 13, 751–754 (2016).

17. Yamamoto, K. et al. Improved virus-induced gene silencing allows discovery of a serpentine synthase gene in *Catharanthus roseus*. Plant Physiol. 187, 846–857 (2021).

18. Mint Evolutionary Genomics Consortium. Phylogenomic Mining of the Mints Reveals Multiple Mechanisms Contributing to the Evolution of Chemical Diversity in Lamiaceae. Mol. Plant 11, 1084–1096 (2018).

19. Lichman, B. R., Godden, G. T. & Buell, C. R. Gene and genome duplications in the evolution of chemodiversity: perspectives from studies of Lamiaceae. Curr. Opin. Plant Biol. 55, 74–83 (2020).

20. Nützmann, H.-W. et al. Active and repressed biosynthetic gene clusters have spatially distinct chromosome states. Proc. Natl. Acad. Sci. U. S. A. 117, 13800–13809 (2020).

21. Xing, H., Wu, Y., Zhang, M. Q. & Chen, Y. Deciphering hierarchical organization of topologically associated domains through change-point testing. BMC Bioinformatics 22, 183 (2021).

22. Rao, S. S. P. et al. A 3D map of the human genome at kilobase resolution reveals principles of chromatin looping. Cell 159, 1665–1680 (2014).

23. Stavrinides, A. et al. Structural investigation of heteroyohimbine alkaloid synthesis reveals active site elements that control stereoselectivity. Nat. Commun. 7, 12116 (2016).

24. Caputi, L. et al. Missing enzymes in the biosynthesis of the anticancer drug vinblastine in Madagascar periwinkle. Science 360, 1235–1239 (2018).

25. Carqueijeiro, I. et al. Two Tabersonine 6,7-Epoxidases Initiate Lochnericine-Derived Alkaloid Biosynthesis in *Catharanthus roseus*. Plant Physiol. 177, 1473–1486 (2018).

26. Laflamme, P., St-Pierre, B. & De Luca, V. Molecular and biochemical analysis of a Madagascar periwinkle root-specific minovincinine-19-hydroxy-O-acetyltransferase. Plant Physiol. 125, 189–198 (2001).

27. St. Pierre, B. & De Luca, V. A cytochrome P-450 monooxygenase catalyzes the first step in the conversion of tabersonine to vindoline in *Catharanthus roseus*. Plant Physiol. 109, 131–139 (1995).

28. Burlat, V., Oudin, A., Courtois, M., Rideau, M. & St-Pierre, B. Co-expression of three MEP pathway genes and geraniol 10-hydroxylase in internal phloem parenchyma of *Catharanthus roseus* implicates multicellular translocation of intermediates during the biosynthesis of monoterpene indole alkaloids and isoprenoid-derived primary metabolites. Plant J. 38, 131–141 (2004).

29. Miettinen, K. et al. The seco-iridoid pathway from *Catharanthus roseus*. Nat. Commun. 5, 3606 (2014).

30. Guirimand et al. Strictosidine activation in Apocynaceae: towards a “nuclear time bomb”? BMC Plant Biol. 10, 182 (2010).

31. Guirimand, G. et al. The subcellular organization of strictosidine biosynthesis in *Catharanthus roseus* epidermis highlights several trans-tonoplast translocations of intermediate metabolites. FEBS J. 278, 749–763 (2011).

32. Simkin, A. J. et al. Characterization of the plastidial geraniol synthase from Madagascar periwinkle which initiates the monoterpenoid branch of the alkaloid pathway in internal phloem associated parenchyma. Phytochemistry 85, 36–43 (2013).

33. Stuart, T. et al. Comprehensive Integration of Single-Cell Data. Cell 177, 1888–1902.e21 (2019).

34. Macosko, E. Z. et al. Highly Parallel Genome-wide Expression Profiling of Individual Cells Using Nanoliter Droplets. Cell 161, 1202–1214 (2015).

35. Di Fiore, S., Hoppmann, V., Fischer, R. & Schillberg, S. Transient gene expression of recombinant terpenoid indole alkaloid enzymes in *Catharanthus roseus* leaves. Plant Mol. Biol. Rep. 22, 15–22 (2004).

36. Zhang, H. et al. The basic helix-loop-helix transcription factor CrMYC2 controls the jasmonate-responsive expression of the ORCA genes that regulate alkaloid biosynthesis in *Catharanthus roseus*. Plant J. 67, 61–71 (2011).

37. Van Moerkercke, A. et al. The basic helix-loop-helix transcription factor BIS2 is essential for monoterpenoid indole alkaloid production in the medicinal plant *Catharanthus roseus*. Plant J. 88, 3–12 (2016).

38. Van Moerkercke, A. et al. The bHLH transcription factor BIS1 controls the iridoid branch of the monoterpenoid indole alkaloid pathway in *Catharanthus roseus*. Proc. Natl. Acad. Sci. U. S. A. 112, 8130–8135 (2015).

39. Peebles, C. A. M., Hughes, E. K., Shanks, J. V. & -Y, S. K. Transcriptional response of the terpenoid indole alkaloid pathway to the overexpression of ORCA3 along with jasmonic acid elicitation of *Catharanthus roseus* hairy roots over time. Metab. Eng. 11, 76–86 (2009).

40. Paul, P. et al. A differentially regulated AP2/ERF transcription factor gene cluster acts downstream of a MAP kinase cascade to modulate terpenoid indole alkaloid biosynthesis in Catharanthus roseus. New Phytol. 213, 1107–1123 (2017).

41. Misra, B. B., Assmann, S. M. & Chen, S. Plant single-cell and single-cell-type metabolomics. Trends Plant Sci. 19, 637–646 (2014).

42. de Souza, L. P., Borghi, M. & Fernie, A. Plant Single-Cell Metabolomics—Challenges and Perspectives. Int. J. Mol. Sci. 21, 8987 (2020).

43. Guo, S., Zhang, C. & Le, A. The limitless applications of single-cell metabolomics. Curr. Opin. Biotechnol. 71, 115–122 (2021).

44. Yamamoto, K. et al. The complexity of intercellular localisation of alkaloids revealed by single-cell metabolomics. New Phytol. 224, 848–859 (2019).

45. Yamamoto, K. et al. Cell-specific localization of alkaloids in *Catharanthus roseus* stem tissue measured with Imaging MS and Single-cell MS. Proc. Natl. Acad. Sci. U. S. A. 113, 3891–3896 (2016).

46. Carqueijeiro, I. et al. Isolation of Cells Specialized in Anticancer Alkaloid Metabolism by Fluorescence-Activated Cell Sorting. Plant Physiol. 171, 2371–2378 (2016).

47. Payne, R. M. et al. An NPF transporter exports a central monoterpene indole alkaloid intermediate from the vacuole. Nat Plants 3, 16208 (2017).

48. Yu, F. & De Luca, V. ATP-binding cassette transporter controls leaf surface secretion of anticancer drug components in *Catharanthus roseus*. Proc. Natl. Acad. Sci. U. S. A. 110, 15830–15835 (2013).

49. Guirimand, G. et al. Spatial organization of the vindoline biosynthetic pathway in *Catharanthus roseus*. J. Plant Physiol. 168, 549–557 (2011).

50. Goodbody, A. E. et al. Enzymic coupling of catharanthine and vindoline to form 3’,4’-anhydrovinblastine by horseradish peroxidase. Planta Med. 54, 136–140 (1988).

51. Costa, M. M. et al. Molecular cloning and characterization of a vacuolar class III peroxidase involved in the metabolism of anticancer alkaloids in *Catharanthus roseus*. Plant Physiol. 146, 403–417 (2008).

52. Denyer, T. et al. Spatiotemporal Developmental Trajectories in the Arabidopsis Root Revealed Using High-Throughput Single-Cell RNA Sequencing. Dev. Cell 48, 840–852.e5 (2019).

53. Ryu, K. H., Huang, L., Kang, H. M. & Schiefelbein, J. Single-Cell RNA Sequencing Resolves Molecular Relationships Among Individual Plant Cells. Plant Physiol. 179, 1444–1456 (2019).

54. Giddings, L.-A. et al. A stereoselective hydroxylation step of alkaloid biosynthesis by a unique cytochrome P450 in *Catharanthus roseus*. J. Biol. Chem. 286, 16751–16757 (2011).

55. Carqueijeiro, I. et al. A BAHD acyltransferase catalyzing 19-O-acetylation of tabersonine derivatives in roots of *Catharanthus roseus* enables combinatorial synthesis of monoterpene indole alkaloids. Plant J. 94, 469–484 (2018).

56. Munkert, J. et al. Iridoid Synthase Activity Is Common among the Plant Progesterone 5beta-Reductase Family. Mol. Plant 8, 136–152 (2014).

57. Smit, S. J. & Lichman, B. R. Plant biosynthetic gene clusters in the context of metabolic evolution. Nat. Prod. Rep. (2022) doi:10.1039/d2np00005a.

58. Ozber, N. & Facchini, P. J. Phloem-specific localization of benzylisoquinoline alkaloid metabolism in opium poppy. J. Plant Physiol. 271, 153641 (2022).

59. Konno, K., Hirayama, C., Yasui, H. & Nakamura, M. Enzymatic activation of oleuropein: a protein crosslinker used as a chemical defense in the privet tree. Proc. Natl. Acad. Sci. U. S. A. 96, 9159–9164 (1999).

60. Calixto, J. B. The role of natural products in modern drug discovery. An. Acad. Bras. Cienc. 91 Suppl 3, e20190105 (2019).

61. Vaillancourt, B. & Buell, C. R. High molecular weight DNA isolation method from diverse plant species for use with Oxford Nanopore sequencing. BioRxiv 783159, (2019).

62. Kolmogorov, M., Yuan, J., Lin, Y. & Pevzner, P. A. Assembly of long, error-prone reads using repeat graphs. Nat. Biotechnol. 37, 540–546 (2019).

63. Vaser, R., Sovic, I., Nagarajan, N. & Sikic, M. Fast and accurate de novo genome assembly from long uncorrected reads. Genome Res. 27, 737–746 (2017).

64. Walker, B. J. et al. Pilon: an integrated tool for comprehensive microbial variant detection and genome assembly improvement. PLoS One 9, e112963 (2014).

65. Durand, N. C. et al. Juicer Provides a One-Click System for Analyzing Loop-Resolution Hi-C Experiments. Cell Systems 3, 95–98 (2016).

66. Dudchenko, O. et al. *De novo* assembly of the *Aedes aegypti* genome using Hi-C yields chromosome-length scaffolds. Science 356, 92–95 (2017).

67. Ludwig, A., Pippel, M., Myers, G. & Hiller, M. DENTIST-using long reads for closing assembly gaps at high accuracy. Gigascience 11, (2022).

68. Flynn, J. M. et al. RepeatModeler2 for automated genomic discovery of transposable element families. Proc. Natl. Acad. Sci. U. S. A. 117, 9451–9457 (2020).

69. Campbell, M. S. et al. MAKER-P: a tool kit for the rapid creation, management, and quality control of plant genome annotations. Plant Physiol. 164, 513–524 (2014).

70. Bao, W., Kojima, K. K. & Kohany, O. Repbase Update, a database of repetitive elements in eukaryotic genomes. Mob. DNA 6, 11 (2015).

71. Chen, N. Using RepeatMasker to identify repetitive elements in genomic sequences. Curr. Protoc. Bioinformatics Chapter 4, Unit 4 10 (2004).

72. Martin, M. Cutadapt removes adapter sequences from high-throughput sequencing reads. EMBnet.journal 17, 10–12 (2011).

73. Kim, D., Paggi, J. M., Park, C., Bennett, C. & Salzberg, S. L. Graph-based genome alignment and genotyping with HISAT2 and HISAT-genotype. Nat. Biotechnol. 37, 907–915 (2019).

74. Kovaka, S. et al. Transcriptome assembly from long-read RNA-seq alignments with StringTie2. Genome Biol. 20, 278 (2019).

75. Li, H. Minimap2: pairwise alignment for nucleotide sequences. Bioinformatics 34, 3094– 3100 (2018).

76. Hoff, K. J., Lomsadze, A., Borodovsky, M. & Stanke, M. Whole-Genome Annotation with BRAKER. in Gene Prediction: Methods and Protocols (ed. Kollmar, M.) 65–95 (Springer New York, 2019).

77. Haas, B. J. et al. Improving the Arabidopsis genome annotation using maximal transcript alignment assemblies. Nucleic Acids Res. 31, 5654–5666 (2003).

78. Lee, E. et al. Web Apollo: a web-based genomic annotation editing platform. Genome Biol. 14, R93 (2013).

79. Lamesch, P. et al. The Arabidopsis Information Resource (TAIR): improved gene annotation and new tools. Nucleic Acids Res. 40, D1202–10 (2012).

80. El-Gebali, S. et al. The Pfam protein families database in 2019. Nucleic Acids Res. 47, D427–D432 (2019).

81. Trapnell, C. et al. Transcript assembly and quantification by RNA-Seq reveals unannotated transcripts and isoform switching during cell differentiation. Nat. Biotechnol. 28, 511–515 (2010).

82. Kaminow, B., Yunusov, D. & Dobin, A. STARsolo: accurate, fast and versatile mapping/quantification of single-cell and single-nucleus RNA-seq data. bioRxiv 2021.05.05.442755 (2021).

83. Emms, D. M. & Kelly, S. OrthoFinder: phylogenetic orthology inference for comparative genomics. Genome Biol. 20, 238 (2019).

84. Lopez-Anido, C. B. et al. Single-cell resolution of lineage trajectories in the Arabidopsis stomatal lineage and developing leaf. Dev. Cell 1043-1055.e4 (2021).

85. Kim, J.-Y. et al. Distinct identities of leaf phloem cells revealed by single cell transcriptomics. Plant Cell 33, 511–530 (2021).

86. Farmer, A., Thibivilliers, S., Ryu, K. H., Schiefelbein, J. & Libault, M. Single-nucleus RNA and ATAC sequencing reveals the impact of chromatin accessibility on gene expression in Arabidopsis roots at the single-cell level. Mol. Plant 14, 372–383 (2021).

87. Csardi, G. & Nepusz, T. The Igraph Software Package for Complex Network Research. InterJournal Complex Systems, 1695 (2005).

88. Smith, C. A., Want, E. J., O’Maille, G., Abagyan, R. & Siuzdak, G. XCMS: processing mass spectrometry data for metabolite profiling using nonlinear peak alignment, matching, and identification. Anal. Chem. 78, 779–787 (2006).

89. Tautenhahn, R., Böttcher, C. & Neumann, S. Highly sensitive feature detection for high resolution LC/MS. BMC Bioinformatics 9, 504 (2008).

90. Kuhl, C., Tautenhahn, R., Böttcher, C., Larson, T. R. & Neumann, S. CAMERA: an integrated strategy for compound spectra extraction and annotation of liquid chromatography/mass spectrometry data sets. Anal. Chem. 84, 283–289 (2012).

91. Berrow, N. S. et al. A versatile ligation-independent cloning method suitable for high-throughput expression screening applications. Nucleic Acids Res. 35, e45 (2007).

